# Rapid evolution of recombination landscapes during the divergence of cichlid ecotypes in Lake Masoko

**DOI:** 10.1101/2024.03.20.585960

**Authors:** Marion Talbi, George F. Turner, Milan Malinsky

## Abstract

Variation of recombination rate along the genome is of crucial importance to rapid adaptation and organismal diversification. Many unknowns remain regarding how and why recombination landscapes evolve in nature. Here, we reconstruct recombination maps based on linkage disequilibrium and use subsampling and simulations to derive a new measure of recombination landscape evolution: the Population Recombination Divergence Index (PRDI). Using PRDI, we show that fine-scale recombination landscapes differ substantially between two cichlid fish ecotypes of *Astatotilapia calliptera* that diverged only ∼2,500 generations ago. Perhaps surprisingly, recombination landscape differences are not driven by divergence in terms of allele frequency (*F*_ST_) and nucleotide diversity (*Δ*(*π*)): although there is some association, we observe positive PRDI in regions where *F*_ST_ and *Δ*(*π*) are zero. We found a stronger association between evolution of recombination and 47 large haplotype blocks that are polymorphic in Lake Masoko, cover 21% of the genome, and appear to include multiple inversions. Among haplotype blocks, there is a strong and clear association between the degree of recombination divergence and differences between ecotypes in heterozygosity, consistent with recombination suppression in heterozygotes. Overall, our work provides a holistic view of changes in population recombination landscapes during early stages of speciation with gene flow.

## Introduction

Meiotic recombination is central to genetics and to evolution in sexually reproducing organisms. It facilitates rapid adaptation by generating new combinations of alleles (Rice and Chippindale 2001; Nielsen 2006; McDonald et al. 2016), but in some contexts can also hinder adaptation by breaking up locally adapted haplotypes (Ortiz-Barrientos et al. 2016; Schluter and Rieseberg 2022). A substantial body of theory has been developed describing genetic variants that regulate recombination rates, so-called ‘recombination modifiers’, and the conditions under which such modifier variants would be selected for or against (Nei 1967; Feldman et al. 1996; Coop and Przeworski 2007). Many genetic variants are known to affect the recombination rates and the positioning of recombination events (Halldorsson et al. 2019; Rowan et al. 2019). For example, recombination suppression is often facilitated by larger structural genetic variants, especially inversions (Jay et al. 2018; Todesco et al. 2020), although insertions, deletions, and sequence translocations have also been implicated (Kent et al. 2017; Rowan et al. 2019; Schluter and Rieseberg 2022). Even single nucleotide polymorphisms (SNPs) can modify recombination rates at specific loci, as in the case of the ade6-M26 mutation which creates a hotspot of 10 to 15x elevated recombination in yeast (Ponticelli et al. 1988; Szankasi et al. 1988).

Recombination is also subject to forces that appear largely decoupled from organismal adaptation or diversification. First, it must fulfil its essential role in meiosis and chromosome segregation (Petronczki et al. 2003). This virtually ubiquitous requirement provides a lower bound of one recombination event per chromosome (Henderson and Bomblies 2021) and contributes to limiting average recombination rates to a relatively narrow range above this minimum via the mechanism of ‘crossover interference’ (Otto and Payseur 2019). Second, in some vertebrate species, principally in mammals, recombination is directed towards binding sites of the zinc-finger protein PRDM9 (Baudat et al. 2010; Myers et al. 2010; Baker et al. 2017; Cavassim et al. 2022). In these species, rapid evolution of recombination landscapes is mediated by intra-genomic conflict (Úbeda and Wilkins 2010; Latrille et al. 2017; Baker et al. 2023) and genetic variants altering recombination landscapes in this way are therefore usually studied through the prism of internal genome dynamics and not within the framework of traditional recombination modifier theory (Genestier et al. 2023).

Outside of mammals, in most other vertebrates, recombination does not appear to be associated with PRDM9 binding sites (Cavassim et al. 2022). Species lacking the PRDM9 mechanism have elevated recombination rates at and around genomic features such as CpG islands and promoters, likely due to the greater chromatin accessibility in these regions (Baker et al. 2017) and there is evidence that species lacking PRDM9 have more conserved recombination landscapes (Lam and Keeney 2015; Singhal et al. 2015). However, the association with these genomic features is only partial (Singhal et al. 2015) and recombination rates do evolve also in species without PRDM9 (Ritz et al. 2017; Samuk et al. 2020; Déserts et al. 2021). It has even been suggested that in stickleback fish, hotspots may evolve at similar rates to those observed in species with PRDM9 (Shanfelter et al. 2019).

In the presence of gene-flow, recombination counteracts the buildup of linkage among genetic loci that contribute to population divergence and, ultimately, speciation (Felsenstein 1981; Barton 2020; Butlin et al. 2021). Consistent with the important role in this context, a recombination landscape, together with natural selection, shape the distribution of genomic regions of divergence along each chromosome (Ravinet et al. 2017; Duranton et al. 2018; Schumer et al. 2018; Martin et al. 2019). The recombination landscape itself is an evolving dynamic parameter (Ritz et al. 2017; Samuk et al. 2020; Déserts et al. 2021), which should be taken into account in genomic studies of speciation (Ortiz-Barrientos et al. 2016; Ortiz-Barrientos and James 2017). In recent years, there has been a growing appreciation of the role of recombination suppression in organismal diversification (Schluter and Rieseberg 2022). These efforts often concentrated on specific large inversions or other low-recombining haplotype blocks (Jay et al. 2018; Faria et al. 2019a; Todesco et al. 2020; Reeve et al. 2023). In addition, a recent study based on genome-wide estimates of recombination rates supports the notion that cis-acting recombination modifiers play an important role in promoting adaptive divergence between populations (Venu et al. 2024). Building upon this initial progress, additional comparisons of recombination landscapes in different species and across different levels of divergence will be required to understand where in the genome, how fast, and by what mechanisms recombination rates evolve and, ultimately, the interplay with natural selection and organismal evolution.

The Lake Masoko system presents a well-suited opportunity to study the evolution of recombination rates in the context of organismal diversification. Lake Masoko is a small (∼670m in diameter) crater lake in Southern Tanzania (**Fig 1A**) and is approximately 50k years old (Barker et al. 2003). Two ecotypes of the cichlid fish species *Astatotilapia calliptera* have evolved within this lake – the shallow-water ‘littoral’ and the deep-water ‘benthic’. They differ from each other in several ecologically important traits and, while almost half of the sites have zero *F*_ST_, there is elevated allele frequency divergence at about a hundred of well demarcated genomic regions – islands of divergence (Malinsky et al. 2015). Several other fish species belonging to the same clade (Percomorpha) lack a functional PRDM9, although *A. calliptera* itself has not been tested (Baker et al. 2017; Cavassim et al. 2022).

**Figure 1:**
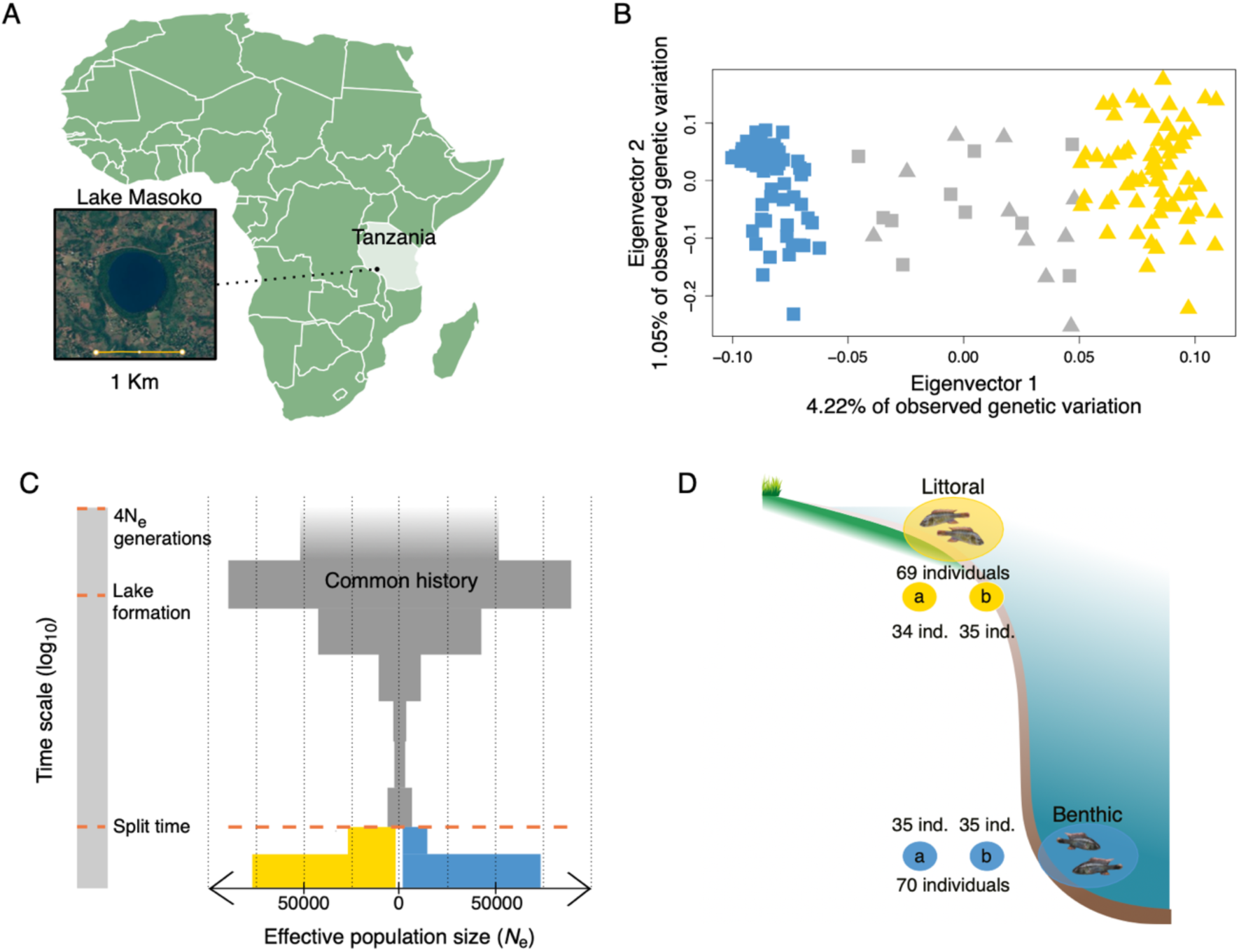
Study system and demographic history. (**A**) Lake Masoko is a circular small (∼670m diameter) maar-type volcanic crater lake located in the East African rift valley in southern Tanzania. (**B**) A principal component analysis based on 784974 SNPs. Some individuals labelled in the field as ‘benthic’ or ‘littoral’ turned out to be genetically admixed. The genetic maps presented and examined in this study are based on individuals with little to no admixture, highlighted in yellow for littoral and blue for benthic. (**C**) SMC++ inference of demographic history – the changes in effective population sizes (*N*e; x axis) – through time (y axis). After the split time, the population sizes for the littoral ecotype are shown in yellow, and for the benthic in blue. (**D**) We divided the individuals from each cichlid ecotype (littoral, benthic) of Lake Masoko into two independent subsets, ‘subset a’ and ‘subset b’. Each subset, and the recombination maps inferred using the subsets, represent biological replicates.

Over the last decade or so, many empirical studies used recombination estimates based on linkage disequilibrium (LD) – patterns of non-random association of genetic variants in a sample of individuals from a population (Auton et al. 2012; Singhal et al. 2015; Shanfelter et al. 2019; Spence and Song 2019). LD-based methods estimate a ‘population recombination rate’ (ρ) – a population genetic parameter which is a product of effective population size (*N*_e_) and the historical per-generation recombination rate averaged over the ancestry of the sampled individuals. Differences between LD-based recombination maps can arise from sampling variance, methodological limitations, different ancestries across samples, and factors affecting *N*_e_ such as demographic change and natural selection (Coop and Przeworski 2007; Peñalba and Wolf 2020; Samuk et al. 2020). Therefore, it is challenging to use LD-based methods to compare recombination landscapes between populations and species to track their evolution.

In this study, we reconstruct genetic maps from patterns of LD in whole genome population genetic data of 70 benthic and 69 littoral individuals to investigate the evolution of recombination landscapes in Lake Masoko. By carefully controlling for confounding factors, we demonstrate that the population recombination landscapes are considerably different despite the recent split time between these ecotypes and we quantify the degree of this population recombination rate evolution. The regions where recombination rates differ significantly are not distributed equally across the genome. We show a link with genetic differentiation, as measured for example by *F*_ST_, and with larger haplotype blocks, although neither of these fully explain the recombination rate divergence. We found a partial copy of PRDM9 in the *A. calliptera* genome. However, its predicted binding sites do not show any relationship with recombination rates, which is consistent with previous studies in fishes with partial PRDM9 (Baker et al. 2017). Overall, our study quantifies and provides new insights into rapid population recombination rate evolution in the context of sympatric ecotype divergence.

## Material and Methods

### Variant calling and filtering

Genomic DNA from a total of 336 individuals from Lake Masoko has been sequenced on the Illumina HiSeq X Ten platform, obtaining 150bp paired end reads (NCBI Short Read Archive, BioProject ID: PRJEB27804). The reads were aligned to the *Astatotilapia calliptera* reference genome fAstCal1.5 (GenBank ID: GCA_900246225.6) using bwa-mem v.0.7.17 (Li 2013), with median depth per-individual of 15.8x (min = 12.4x, max = 22.1x). The reference sequence is based on an *A. calliptera* sample from the Itupi stream which is a close outgroup to Lake Masoko. We used the `MarkDuplicates` tool from the Picard package v.2.26.6 to tag PCR and optical duplicate reads and GATK v.4.2.3 (DePristo et al. 2011) to call variants, using `HaplotypeCaller` in GVCF mode for each individual separately followed by joint genotyping using `GenotypeGVCFs` with the `--include-non-variant-sites` option.

Next, we generated a *callability mask* to identify and filter out the regions of the genome where we were unable to confidently call variants. The mask included: i) sites with an overall read depth cutoffs based on examining a depth histogram (< 3800 or > 5700); ii) sites where more than 10% of individuals had missing genotypes; iii) sites identified by GATK as low quality (with the `LowQual` tag) and iv) sites with poor mappability. Specifically, to obtain the mappability information, we broke down the genome into overlapping k-mers of 150bp (matching the read length), mapped these k-mers back to the genome, and masked all sites where fewer than 90% of k-mers mapped back to their original location perfectly and uniquely. In total, the callability mask comprised of 311 million bp, or about 35% of the genome. In addition to applying the callability mask, we used several hard filters based on GATK best practices, specifically `MQ<40`, `FS>40`, `QD<2` and `ExcessHet>40`. These additional filters removed fewer than 1% of the remaining SNPs. After discarding 1.4 million of multiallelic sites and indels, the filtered VCF contained 3.86 million SNPs.

### Sample selection for recombination analyses

We used the full set of 336 available individuals for variant calling because the inclusion of more samples leads to more accurate genotyping. However, in this study we were specifically interested in differences between the littoral and benthic ecotypes of Lake Masoko. Therefore, we retained 80 individuals assigned in the field as benthic and 79 assigned as littoral and excluded 201 other individuals which were not assigned to either ecotype because they were juveniles, females (neither category show the ecotype-distinct male breeding colors), or putative hybrids. To check the validity of these field assignments, we first built a neighbor-joining tree based on a genetic distance matrix, i.e., the average number of single-nucleotide differences between each pair of individuals, using the `stats` command from the evo package v.0.1 r28 and the `--diff-matrix` option. The pairwise difference matrix was divided by the callable genome size to obtain pairwise distances per base pair and this was then used as input into the `nj()` tree-building function implemented in the package *ape* in R (Paradis et al. 2004). Next, for principal component analysis we used smartPCA (Patterson et al. 2006) on data filtered for minor allele frequency >=0.05 using plink v1.9 (Purcell et al. 2007) with the `--maf 0.05` option and LD pruned using the plugin `+prune` from bcftools v.1.16 (Danecek et al. 2021) with the `-m 0.8 -w 1000` options. After we identified and removed 20 genetically intermediate samples (see **Figure 1C** and **Supplementary Figure 1**), the final VCF with 139 individuals was composed of 3.3 million biallelic SNPs.

### Genome annotation

Because the fAstCal1.5 assembly was not annotated by NCBI, we ‘lifted over’ the annotation from the older fAstCal1.2 assembly (GenBank: GCA_900246225.3). We used the UCSC paradigm (Miller et al. 2007) to generate a pairwise whole genome alignment between fAstCal1.2 and fAstCal1.5 assembly. Afterwards, we used the UCSC `liftOver` tool to translate the NCBI Annotation Release 100 to the new coordinates.

### Inference of demographic history and estimation of the level of gene flow

To estimate split time between the ecotypes and changes in effective population size (*N*_*e*_) through time, we used smc++ v.1.15.4 (Terhorst et al. 2017), using the sequence of smc++ commands: `vcf2smc`-> `estimate`-> `split`, To translate the time axis into number of generations, we used the cichlid-specific mutation rate estimate of *μ* = 3.5 x10^’#^ per bp per generation with 95% confidence interval (1.6 x10 ^‘#^, 4.6 x10^‘#^) (Malinsky et al. 2018). Next, we used fastsimcoal2.7 (Excoffier et al. 2021) to estimate the level of gene flow between the two ecotypes. In fastsimcoal, we entered the split time and changes in *N*_*e*_ as inferred by smc++ as fixed parameters and estimated continuous asymmetrical migration rates after the ecotype split. We ran 30 simulations with different starting parameter values, which revealed two local peaks in the likelihood surface (**Supplementary Figure 2**). To reduce the confounding effects of selection in these demographic analyses we used only sites from non-coding regions of the genome, masking all annotated exons, introns, and promoters.

### Subsampling and bootstrap

We first used the `shuf-n` command to randomly draw a set of 35 individuals from each ecotype to for the first subset. The remaining individuals (35 littoral and 34 benthic) then formed the second subset. Therefore, these (a) and (b) subsets (**Figure 1B**) are independent, in the sense that they are composed of non-overlapping sets of individuals. We then generated a separate VCF file for each subset using the bcftools v.1.16 `view` command and used these VCFs for recombination map reconstruction. We repeated this random sampling procedure (and the following genetic map reconstruction) to obtain nine bootstrap replicates over individuals.

### Inference of recombination rates

We used the `pyrho` software (Spence and Song 2019) to infer recombination rates along the genome based on patterns of linkage disequilibrium. We choose `pyrho` because it accounts for demography, i.e., changes in *N*_*e*_through time, and because its performance does not depend on haplotype phasing – it performs equally well with phased and unphased data [as described in fig. S8 of (Spence and Song 2019) and confirmed by our own simulations (data not shown)].

Specifically, first, to build likelihood tables between each biallelic sites, we used the `make_table` command with *μ* = 3.5 × 10^‘#^, demographic history for each of the ecotypes as inferred by smc++, and the Moran approximation with the `–approx` and `–moran_pop_size N` flags where N equals 1.5x the number of haplotypes in each subset. To determine the best parameter to use for the inference of recombination rates, we processed a set of simulation with evolutionary parameters corresponding to the one of our cichlid species (e.g., *µ*, sample size, *N*_*e*_) and chose the value of block penalty that was minimizing the quantity of false negative and false positive with best results obtained for block penalty of 15 and a window size of 50 SNPs. These parameters were then used in all runs of the `optimize` command to infer the recombination maps. The output of `pyrho` contained estimates of recombination rate between each pair of SNPs.

### Neutral coalescent simulations

We used msprime v.1.0.2 (Baumdicker et al. 2021) to simulate genetic data matching the population and demographic histories (split time, *N*_*e*_, and gene flow) that we inferred from empirical data as described above. Because recombination landscapes were the same for both simulated populations and natural selection absent in these simulations, the results from analyzing the simulated data allowed us to better evaluate and interpret the empirical results. We ran 23 simulations – one for each chromosome – using *μ* = 3.5 × 10^-9^ and the empirical recombination maps as input. From each simulation, we sampled 70 individuals from each population, labelled them as ‘benthic’ and ‘littoral’, randomly subsampled the (a) and (b) subsets and further processed the VCF output in the same way as we did for empirical data.

To confirm that the conclusions of this manuscript are robust to the specifics of demographic inference, we ran the simulations with an extended set of split time values (1 000, 2 500, 5 000 and 10 000 generations ago) and with migration rates corresponding to the two local likelihood peaks in fastsimcoal2 inference (‘low migration rates’: 11.5 × 10^‘$^ for littoral to benthic and 7.01 × 10^‘$^ for benthic to littoral; and ‘high migration rates’: 0.0339 for littoral to benthic and 0.0368 for benthic to littoral).

We also used msprime to estimate how the inferred recombination maps reflect the relative contributions of recombination events that happened in the common history of the ecotypes vs. events that happened after their split. To do this, we counted the the number of local genealogies, which reflects the recombination events that changed the local genealogy of the sample. We used the `end_time` option in the `sim_ancestry()` function of msprime to stop the simulation at the split time and counted the distinct genealogies at that time point in benthic (*N*_*tb*_) and in littoral (*N*_*tl*_) ecotypes – these counts reflect the genealogy-changing events that happened in the ecotypes after their split. Then we continued the simulation backward in time all the way to the common ancestor of all samples and counted the total number of genealogies (*N*_*t*_). Finally, we calculated the proportion of genealogy-changing events that happened in the ecotypes after their split (*P*_*as*_) as:

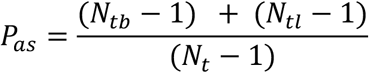

The scripts used to run these simulations are available from GitHub.

### Processing and comparisons of recombination maps

We used the `PhysicalWindowAverages` command from the evo package v.0.1 r28 to obtain mean recombination rates in identical 2Kb windows for all datasets, which facilitated easy comparisons between different maps. The correlations, map distances, and other comparisons were then calculated using R scripts, available from GitHub.

The first set of comparisons among recombination maps was based on correlations, which has allowed us to define a measure of divergence which we call the Population Recombination Divergence Index (PRDI). Briefly, we calculated pairwise Spearman correlations of recombination maps in 2kb windows, binned per 5Mb regions along the genome and obtained the median of these correlations. We first calculated the correlations for the within-ecotype replicate maps (medians denoted *m*_wb_ and *m*_wl_ for within-benthic and within-littoral replicates respectively). Then we calculated correlations for map comparisons between ecotypes, denoting the median of all these comparisons *m*_b_. Between-ecotype correlations are generally lower than correlations for within-ecotype replicates and we measure this difference using the medians as:

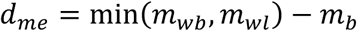

The d_*me*_measure is derived from the empirical data. We also define an analogous measure, d_*m*s_ based on the simulated data to account for the separation of recent benthic vs. littoral ancestry. Finally, PRDI, our measure of recombination landscape divergence is defined as the difference between these two values:

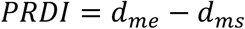

The second set or comparisons was based on recombination map distances. Map distances were calculated in non-overlapping 100kb windows (i.e. vectors of 50 values for 2kb each) using the `dist()` function in R. We use *log_10_* transformed absolute (Manhattan) distances, because these are straightforward to interpret: a *log_10_* distance of 1 signifies an average difference in recombination estimates of one order of magnitude. To find the regions of the genome where the recombination distance between the ecotypes was significantly elevated, which we refer to as *Δ*(*r*) outliers, we used the following procedure. We first calculated the recombination distances in comparisons of within-ecotype replicate maps (denoted *Δ*(*r*)_*w*). Then we calculated the analogous measure for map comparisons between ecotypes (denoted *Δ*(*r*)_b). Finally, we calculated the standard deviation of the *log_10_* transformed *Δ*(*r*)_*w* measure across the bootstrap replicates (we denote this standard deviation as sd_*_). We then refer to any 100kb interval of the genome as a *Δ*(*r*) outlier if it satisfies the following inequality:

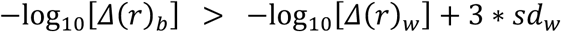

### Filtered maps

The reliability of LD-based recombination rate inference varies across the genome, depending on several factors, including miscalled variants, errors in the reference genome, and amount of genetic variation (i.e., amount of data available for inference). To understand how our results are affected by these factors, we generated filtered recombination maps where the less reliable regions of the genome were masked. While the main figures of this manuscript report the results for the raw maps, we conducted many of the key analyses also using the filtered maps and present these results as supplementary figures.

First, errors in the reference genome can mistakenly place in physical proximity genetic variants that have large genetic distances between them. Therefore, we masked intervals 50kb upstream and 50kb downstream of each joint between contiguous sequences (contig joint) in the assembly. There were 514 joints on the 22 chromosomes, leading to masking of 50.46Mb of sequence. Second, we used the callability mask produced for variant filtering (see above). A lack of data can make recombination inference difficult. We reduced this effect by masking each 100kb region within which more than 70% were not callable. Third, to exclude regions where the recombination maps showed especially elevated sampling noise, we took advantage of the bootstrap runs and masked all 100kb regions where the difference in inferred recombination rates between bootstrap runs was greater than one order of magnitude. Overall, the filtered recombination maps had masked a total of 264 Mb, or approximately 30.6% of the chromosomes, which is about 5% less than the was filtered out by the callability mask during in genotype filtering.

### Hotspot analyses

A recombination hotspot is a narrow region of unusually elevated recombination rate. When searching for hotspots in our data, we required the local recombination rate estimate in any inter-SNPs interval to be at least five times higher than a background rate. For the background recombination rate, we applied three definitions the: (i) mean rate in 40kb around the interval (20kb before and 20kb after), (ii) mean rate in 1Mb around the interval, and (iii) mean recombination rate for the whole chromosome as in (Halldorsson et al. 2019). In most cases, several neighboring intervals were identified as being a part of a hotspot and these intervals were merged using the bedtools v2.29.2 `merge` command if the distance between such intervals was less than 1kb. Proportion of overlap between hotspots from different maps were calculated using the `intersect` command from bedtools v2.29.2 with default parameters, meaning that hotspots were considered overlapping if they shared at least 1bp. When considering the mean recombination rates around hotspots (**Supplementary Figure 5B**), we normalized the highest point in each hotspot to equal 1.0, so that all hotspots were considered equal. Some hotspots were very long and contained implausibly large fractions of recombination, a phenomenon also reported by other studies (Auton et al. 2012; Hoge et al. 2023). We removed hotspots longer than 5kb from the above analyses and from the search for sequence motifs described below.

### Measures of genetic differentiation

To assess the degree of genetic differentiation between the benthic and littoral ecotypes for windows along the genome we calculated *F*_ST_ and the difference in nucleotide diversity (*π*), which we call *Δ*(*π*). Our *F*_ST_ calculation implements the Hudson estimator, as defined in equation 10 in (Bhatia et al. 2013), using ‘ratio of averages’ to combine estimates across multiple variants. To calculate nucleotide diversity for each ecotype, we divide the average number of differences between any two haplotypes by the number of callable sites in each genomic window. These calculations are implemented in the `Fst` command of the evogenSuite software, with the `--accessibleGenomeBED` option providing an inverse of the callability mask. We did this (i) for physical windows of 2Kb (`-f` option) and (ii) for 20 SNPs windows along the genome (`-w` option).

### Distance from CpG islands and transcription start sites

We used two different definitions of CpG islands (GpGi). First, consistent with (Baker et al. 2017), we used the the `maskOutFa` and `cpg_lh` command from UCSC utils. This approach stipulates a minimum of 50% GC content for defining a CpGi. This resulted in 17 000 CpG islands representing a total of 8.5 Mb. Because GC content in the genome is lower in percomorpha that in vertebrates due to less biased gene conversion toward GC (Escobar et al. 2011) and constraining the definition of CpG island by GC content may not be the most appropriate here. We thus also used the `cpgplot-minoe 0.6-minpc 0` command from `EMBOSS` software (Rice et al. 2000), which resulted in a much greater number (228 069) of CpGi. For the Transcription Starting Sites (TSS), we used the genome annotation described before. We then used the `intersect-v` and `closest` command from `bcftools v.2` to obtain the mean recombination rates in 2Kb and 10Kb associated with the distance of the closest TSS non overlapping with a CpGi and vice versa. We finally used the `loess` function in R to obtain the relative recombination rate depending on the distance with the closest TSS or CpGi.

### Inference of haplotype blocks

To discover large-scale variation shared by loci along the genome, we used the program `lostruct` that visualizes the local effect of population structure (Li and Ralph 2018). `lostruct` summarize the pattern of relatedness in a local PCA for nonoverlapping windows along the genome and calculate the dissimilarities between each pair of local PCAs. It then uses multidimensional scaling (MDS) to visualize relationships between windows. We ran `lostruct` for each chromosome separately on 100 SNPs windows. We then plot the first and second axis of the MDS against the genome position. We manually identified 47 regions with high MDS values and high difference in the MDS scores (**Supplementary Figure 10**). To visualize population structure in `lostruct` outliers, we used `smartPCA` (Patterson et al. 2006) on data filtered for minor allele frequency >=0.05 using plink v1.9 (Purcell et al. 2007) with the `--maf 0.05` option but did not filter based on LD.

### Characterization of haplotype blocks

To better understand the evolutionary history of each of the 47 haplotype blocks we calculated and looked at the relationships of a range of population genetic statistics in these regions. First, to test if the proportion of high-heterozygosity individuals was different between ecotypes, we calculated the proportion of heterozygous sites for each individual (H_ind_), used the `kmeans` function in R with `k=2` to assign these values into two clusters, and then used a binomial distribution to quantify the difference in the number of individuals from each ecotype in the cluster with higher H_ind_. (**Figure 4C**; **Supplementary Table 2**). The inbreeding coefficient *F* was calculated for each SNP using our R code. To estimate the time to the most recent common ancestor (TMRCA) we calculated pairwise d_xy_ (i.e. the number of differences in the nucleotide sequence) among all individuals. We took the maximum value of pairwise d_xy_ within each of the 47 local PCA outliers to approximate the TMRCA as 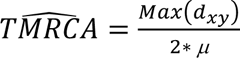 (acknowledging that this can be a slight underestimate of the true TMRCA of individual sequences because of using unphased data).

We then manually assigned ten control regions of a length of 3.9Mb, corresponding to the mean length of the local PCA outlier regions, that we chose to lie in regions of low MDS1 and MDS2 scores. The relative enrichment for each negative value of *F* in the local PCA outliers in comparison with the control regions was calculated as the relative proportion of SNPs from each category (outliers vs. controls) in each interval of *F* values using our R code.

The highly differentiated regions (HDRs) were defined analogously to ref. (Malinsky et al. 2015). We took the top 1% of the *F*_ST_ values in 20 SNP windows (calculated as described above) and merged the windows that were within 10kb of each other. This resulted in 352 HDRs which is very similar to the 344 HDRs found in (Malinsky et al. 2015). The permutation test used to assess significance of mean *F*_ST_ per each local PCA outlier was implemented using custom R script. Briefly, for each local PCA outlier we sampled 1,000 random genomic windows of the same size, obtaining a null distribution of *F*_ST_ values, and considered the *F*_ST_ significantly elevated when it fell within the 5% of this distribution.

### PRDM9 ortholog research and distance from Zinc-Finger binding motif

To find the PRDM9 ortholog in *A. calliptera*, we used the `blastp` command (Altschul et al. 1990) using as query the PRDM9 protein sequence from the Atlantic salmon *Salmo salar* (Gene ID 100380788) against the NCBI RefSeq database (O’Leary et al. 2016) for the species *Astatotilapia calliptera*. Using the best matching protein sequence (XP_026034002.1), we applied a Zinc-Finger (ZF) motif predictor (Persikov et al. 2009; Persikov and Singh 2014) to obtain the position weight matrix representing the binding site prediction based on 11 identified zinc finger domains. We then used the `FIMO` command from the `MEME Suite` (Bailey et al. 2015) on the reference genome of our species, to localize the binding sites. We then used the `intersect-v` and `closest` command from `bcftools v.2` to obtain the mean recombination rates in 2Kb with the distance of the closest ZF binding DNA motif.

### Contribution of genomic features to recombination divergence

In **Table 1**, we summarize how recombination outliers (*Δ*(*r*) outliers) coincide with different genomic regions (10 % regions of higher *F*_ST_, high *Δ*(*π*), `lostruct` outliers, and predicted PRDM9 binding sites). The excess of *Δ*(*r*) outliers in these regions was calculated by dividing the proportion of *Δ*(*r*) outlier sequence length overlapping these regions by the proportion of the genome taken up by the regions. The significance of excess overlap was calculated using 1,000 permutations with the R package `regioneR v. 1.34.0` (Gel et al. 2016).

**Table 1:** Genomic regions underlying recombination rate evolution.

| *Feature* | $\Delta(r)$ outlier<br>proportion | $\Delta(r)$ outlier<br>excess | Permutation<br>p-value | Mean $\Delta(r)$<br>(log10) |
| --- | --- | --- | --- | --- |
| ***High $F_{ST}$ (top 10%)*** | 13.7% | +58.3% | < 0.001 | 0.109 |
| ***High $\Delta(\pi)$ (top 10%)*** | 12.8% | +47.8% | < 0.001 | 0.153 |
| ***Local PCA outliers*** | 31.6% | +47.5% | 0.031 | 0.133 |
| ***PRDM9 binding sites*** | 0.08% | -8.31% | 1 | 0.106 |

## Results

### Study system and demographic history

We obtained whole genome short read sequences of 159 male individuals, each of which was assigned to either the benthic or littoral ecotype based on a field photograph. The sequencing coverage was ∼15x and, after variant calling and filtering, we obtained 3.9 million SNPs that were used for all the following analyses. To check the validity of the field assignment we used the genetic data to run a principal component analysis (PCA; **Figure 1B**) and reconstructed a Neighbor-Joining tree (**Supplementary Figure 1**). While no individuals were misassigned, we identified 20 individuals that appeared to be genetically intermediate which may be the result of recent hybridization. Because our goal was to focus on the differences between the ecotypes, we removed the intermediate individuals (gray in **Figure 1B**) from further analyses.

To obtain a more accurate understanding of the historical demographic context of the ecotype divergence, we first used the program SMC++ (Terhorst et al. 2017) to infer the recent changes in effective population size. Consistent with previous results (Malinsky et al. 2015), we find a bottleneck which may be related to the lake colonization, followed by recent demographic expansions in both ecotypes (**Figure 1C**). This approach also allowed us to re-estimate the split time between the two ecotypes, which we now put at ∼2,500 generations ago (95% confidence interval: 1902 to 5469 generations). While this is considerably older than reported previously (Malinsky et al. 2015), this difference is primarily a result of using the cichlid-specific mutation rate from (Malinsky et al. 2018) in place of the human mutation rate used in the previous study. We also estimated the amount of gene flow between the ecotypes using fastsimcoal2, with best migration rate estimates being 11.5 × 10^-5^ for littoral to benthic and 7.01 × 10^-5^ for benthic to littoral, although higher migration rates of 0.0339 for littoral to benthic and 0.0368 for benthic to littoral had almost equivalent likelihoods and appear more realistic given the number of intermediate individuals we found (**Supplementary Figure 2**; Materials and Methods).

### Population recombination landscapes differ between ecotypes

First, to quantify the effects of sampling variability and methodological limitations, we divided the individuals from each ecotype into two independent subsets and reconstructed a separate recombination map for each subset (**Figure 1D**). Thus, we obtained a total of two replicate maps for each ecotype. Spearman correlation between the replicate maps from the same ecotype (within-ecotype) was 0.77 for within-littoral and 0.71 for within-benthic comparisons at 2kb scale (**Figure 2A**; Materials and Methods). The relatively low correlation coefficients for the within-ecotype replicates reflect a sensitivity of recombination rate inference to sampling variance. Next, we made recombination landscape comparisons between ecotypes (in gray in **Figure 2A**) and found that the correlation coefficients were considerably lower than within ecotypes (**Figure 2A**; mean Spearman correlation = 0.57). The difference in median correlations (within vs. between ecotypes) which we denote d_me_ was thus 0.71 – 0.57 = 0.14 (see Materials and Methods). The key result, *i.e.* that between-ecotype correlations are consistently and considerably lower than for within-ecotype replicates, holds across genomic scales from 2kb to 5Mb (**Supplementary Figure 3**). It is also highly consistent across bootstrap replicates obtained by splitting the ecotypes into different subsamples of individuals (**Supplementary Figure 3**).

**Figure 2:**
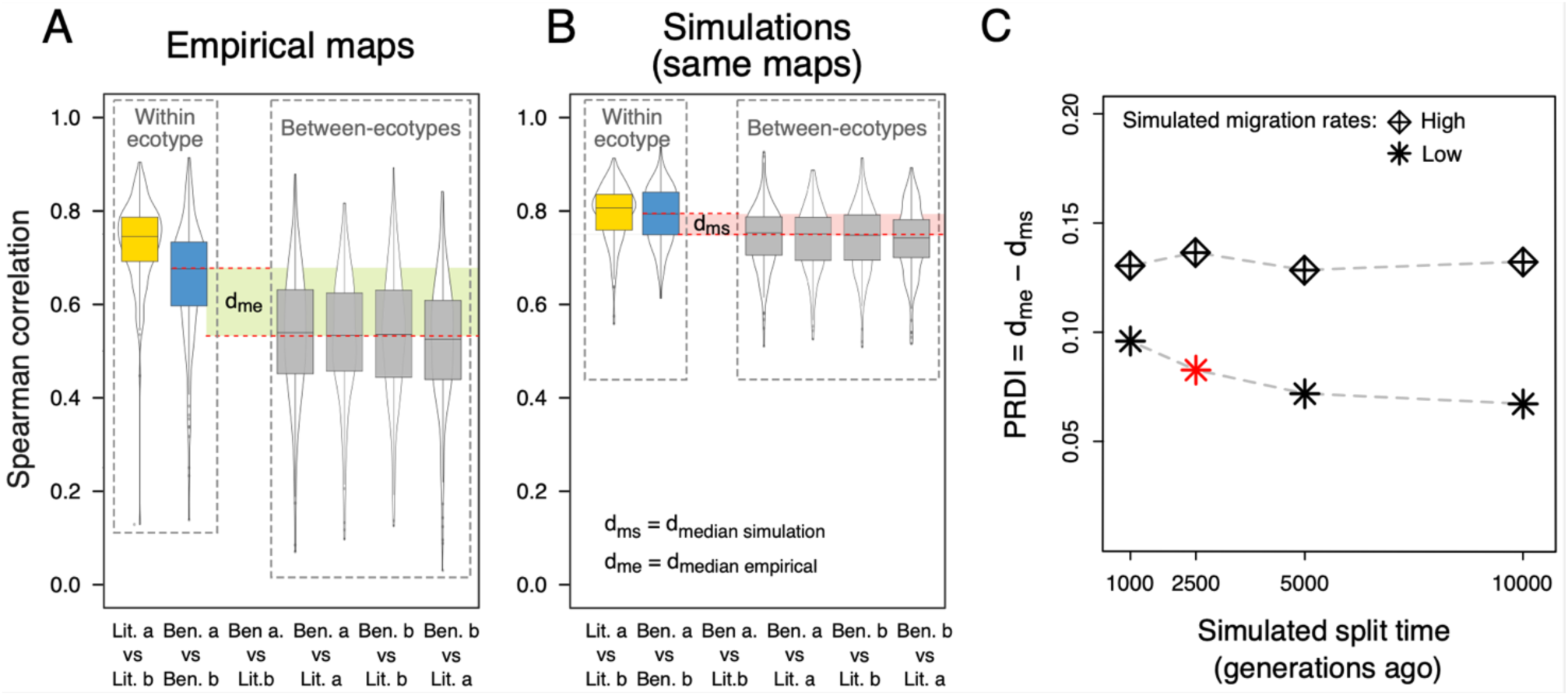
Rapid evolution of the recombination landscapes between the ecotypes. (**A**) Spearman correlation between recombination maps on 2kb scale with each datapoint representing a 5Mb genomic interval. The correlations in between-ecotype comparisons are substantially lower than in the within-ecotype replicate comparisons, with the difference measured by dme (Material and Methods). Blue and yellow colors refer to the benthic and littoral ecotypes respectively. (**B**) Correlations among recombination maps inferred from population genetic data simulated under our best-estimate parameters, matching the demographic history of Lake Masoko. The between-ecotype comparisons also show lower correlations than within-ecotype replicates, reflecting the recent separate ancestries of the ecotypes, but this difference (dms) is lower than dme. (**C**) A measure of recombination landscape evolution, the Population Recombination Divergence Index (PRDI) as a function of different split times and migration rates used for the simulations. Across the entire parameter space, PRDI varies between 0.07 and 0.14. The datapoint highlighted in red is based on the best-estimate parameters shown in (B).

To account for the separation of recent benthic vs. littoral ancestry, which leads to non-random sampling of recombination histories, we conducted neutral coalescent simulations assuming that recombination landscapes were the same for both ecotypes. First, we used msprime to simulate genetic data matching our best estimates of population and demographic histories (split time, *N*_*e*_, and gene flow) that we inferred from empirical data as described above. Based on data simulated under these best-estimate parameters, the within-ecotype vs. between ecotype difference in median recombination map correlations, which we denote d_ms_ was 0.046, was approximately three times lower than what we observed in empirical data (empirical d_me_ = 0.140 vs. simulation d_ms_ = 0.046; **Figure 2B**). We consider this difference (d_me_ - d_ms_) to be a meaningful measure of recombination landscape evolution and refer to it as the Population Recombination Divergence Index (PRDI) (Materials and Methods; **Figure 2C**).

Because the PRDI estimate depends on the parameters used in the simulations, and because there are uncertainties regarding the correct values for these parameters, we conducted additional simulations across a broad range of split times and using both the higher and lower migration rate estimates from fastsimcoal2. We found that with the higher migration rates, PRDI estimates were consistently high, around 0.14 across the full range of split times, because d_ms_ hovered around zero reflecting the intertwined ancestries of the ecotypes in the presence of high migration (**Figure 2C**). With the low migration rates, PRDI was lower and depended on the simulated split time, reaching a minimum of 0.07 when we simulated ecotype divergence 10,000 generations ago (**Figure 2C**). Overall, these results provide evidence that the population recombination landscapes differ between the ecotypes, with the magnitude of divergence estimated by PRDI to be between 0.07 and 0.14.

To further confirm that the observed recombination differences between benthic and littoral were not driven by technical artifacts in regions of the genome where inference is especially error prone, we applied a stringent filtering mask. This mask was based on the: (i) location of contig joins in the assembly, (ii) ability to confidently call SNPs across genomic windows, and (iii) consistency of inferred genetic maps across bootstrap replicates (Material and Methods). In total, we masked 30.6% of the sequence, eliminating a substantial proportion of noise from the inferred recombination maps across all genomic scales from 2kb to 5Mb (**Supplementary Figure 3**). We repeated all the analyses above using these filtered maps and found that, despite the stricness of the filtering, the results, and specifically the d_me_ and PRDI estimates were virtually the same as for the raw maps (**Supplementary Figures 3, 4A**). We also verified that the d_ms_ estimates and PRDI do not depend on the reference map that is used as input for the coalescent simulations (**Supplementary Figure 4**).

In Lake Masoko *A. calliptera*, the ancestry for the sampled individuals extends substantially beyond the ecotype split time (**Figure 1C**), which means that the inferred recombination maps for each ecotype can be interpreted as a mixture of two time periods: (i) recombination events that happened in the common history of the ecotypes and (ii) events that happened after their split. To estimate the contribution of each of these epochs, we used coalescent simulations and found that, across ten simulations with the best-estimate split time of ∼2,500 generations ago, an average 34.9% of recombination events that changed the genealogy of the sample occured after the split (min = 34.4%; max = 35.7%; Materials and Methods). Thsese are the recombination events that make up the differences between the recombination landscapes between the ecotypes.

Recombination landscapes are highly heterogenous, and a large proportion of events tends to occur in so-called ‘hotspots’ (Coop and Przeworski 2007; Peñalba and Wolf 2020). We quantified the heterogeneity of recombination along the genome in the *A. calliptera* of Lake Masoko and found that 50% of all events were concentrated in less than 9.6% of the genome (**Supplementary Figure 5A**; Between 8.9% and 10.4% depending on the ecotype and the subsample). Therefore, the concentration of recombination in hotspots can be considered intermediate - substantially lower than for example in humans, but higher than for example in the plant Arabidopsis thaliana (**Supplementary Figure 5A**). Using a definition of hotspots as having at least 5x higher recombination rate than the 500kb of surrounding sequence, we found on average 2322 hotspots in each recombination map (between 2275 and 2345; **Supplementary Table 1**). Only 41.5 % of hotspots were shared when comparing ‘a’ and ‘b’ replicate maps within the same ecotype, showing that hotspot detection is particularly sensitive to sampling variance. Nevertheless, we again observed the same pattern as for correlations – the comparisions between benthic and littoral maps showed even less hotspot overlap than expected based on simulations (**Supplementary Figure 5**). Qualitatively similar results were obtained using different hotspot definitions (**Supplementary Figure 5**).

### Characterizing recombination rate evolution

The changes in recombination landscapes measured by LD-based maps are based on effective recombination – the recombination events present in the ancestry of the sampled individuals – and are directly influenced by divergent selection between the ecotypes. Therefore, we next investigated the relationship between recombination rates and genetic divergence between the ecotypes in terms of allele frequencies (*F*_ST_) and levels of nucleotide diversity (*Δ*(*π*)) along the genome. As expected, we found that PRDI generally increased with *F*_ST_ and *Δ*(*π*), consitent with an effect of divergent selection on genotypes (**Figure 3A**). However, we observed evolution of recombination landscapes (positive PRDI) also at low levels of genetic divergence and were not limited to regions of particularly high *F*_ST_ or *Δ*(*π*) (**Figure 3A**). Therefore, the evolution of recombination landscapes is not driven by selection on genotypes (e.g. selective sweeps), and/or the effect that differences in *π* can have on the accuracy of recombination inference (**Supplementary Figure 6**; Raynaud et al, 2023). Moreover, equivalent results were obtained for filtered maps, providing additional confidence regarding the robustness of these conclusions (**Supplementary Figure 7**).

**Figure 3:**
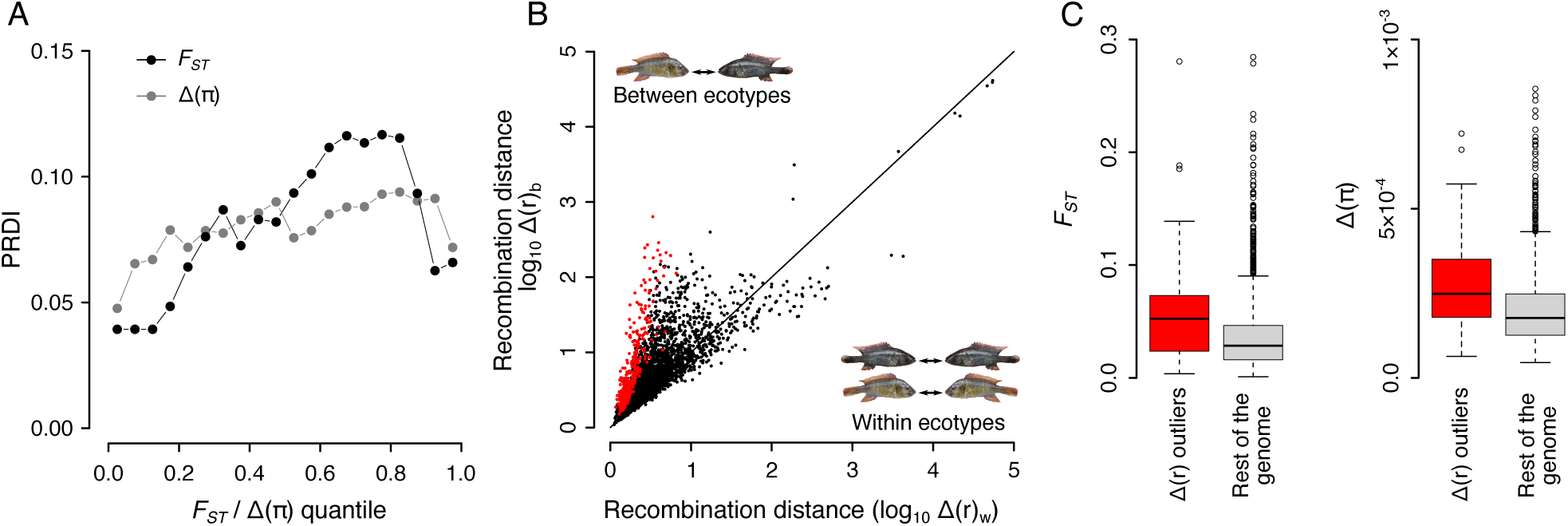
Interplay between recombination and genomic differentiation. (**A**) The Population Recombination Divergence Index (PRDI)_as a function of benthic-littoral *F*ST and of *Δ*(*π*). (**B**) A scatterplot of average recombination map distances in 100kb windows, within biological replicates - *Δ*(*r*)_w_and between the ecotypes - *Δ*(*r*)_b_. Datapoints corresponding to ‘*Δ*(*r*) outliers’ are highlighted in red color. For details see text. (**C**) Comparing the distributions of benthic-littoral *F*ST values and of *Δ*(*π*) within and outside of *Δ*(*r*) outliers.

To explore how the differences in recombination between the ecotypes are distributed across the genome, we calculated the mean difference in recombination rates between the inferred genetic maps in 100kb windows (see **Methods**). The average recombination distance beween benthic and littoral maps (denoted Δ(*r*)_b_) was greater than the distance for within-ecotype replicates (denoted Δ(*r*)_*_) in 83.1% of the windows (**Figure 3B**). This metric allowed us to identify genomic regions with rapidly diverging recombination rates. In the following, we use the ‘net recombination distance’ Δ(*r*) = Δ(*r*)_b_ − Δ(*r*)_*_ and give particular focus to ‘Δ(*r*) outliers’ – regions where the between-ecotype distance is more than three standard deviations higher than within ecotypes. These outliers correspond to 42.7 Mb of sequence, which is about 5% of the genome.

We found that Δ(*r*) outliers are not uniformly distributed across chromosomes; for example, they cover only 1.2% of chromosome 15 (LS420033.2) but over 11.9% of chromosome 1 (LS420019.2) (**Supplementary Figure 8A**). Furthermore, the proportion of outliers across chromosomes is positively correlated with average per-chromosome *F*_ST_. Although this chromosome-wide link is only moderately strong and not statistically significant (Pearson correlation = 0.18; p = 0.42; **Supplementary Figure 8B**), when looking directly at Δ(*r*) outliers, we found that both *F*_ST_ and *Δ*(*π*) were significantly elevated (Mann–Whitney U test: p = 1.88 × 10^‘6^ for *F*_ST_ and p = 5.5 × 10^-5^ for *Δ*(*π*)), clearly confirming that there is an association between allele frequency divergence and the most rapidly evolving population recombination landscapes (**Figure 3C**).

Given the rapid evolution of recombination rates across the genome, we wanted to verify whether the PRDM9 mechanism may be active in Lake Masoko *A. calliptera*. As in several other percomorph species (Cavassim et al. 2022), we found one incomplete PRDM9 ortholog missing the KRAB and SSXRD domains that appear to be necessary for PRDM9 to direct recombination (Baker et al. 2017). Consistent with this, recombination rates were elevated at and near CpG islands (∼1.2x higher; **Supplementary Figure 9**) and transcription start sites (TSS; ∼1.3x higher; **Supplementary Figure 9**), a pattern that is similar to that reported previously for swordtail fish (Baker et al. 2017). Because the PRDM9 zinc finger array in *A. calliptera* was intact, we predicted its binding sites across the genome and found no increase in recombination rates at or near the binding sites (**Supplementary Figure 9**). Overall, these results confirm that PRDM9 does not direct recombination in *A. calliptera* and, therefore, cannot contribute to the rapid evolution of recombination rates in this system.

### Large haplotype blocks contribute to evolution of recombination rates

Ecotype divergence and speciation in the face of gene-flow are often facilitated by regions of suppressed recombination, which allow a buildup of linkage between multiple loci under divergent selection (Faria et al. 2019b). Non-recombining haplotype blocks can be revealed as extended regions of the genome with distinct population structure, substantially different from the genome-wide average (Ma and Amos 2018; Mérot 2020; Todesco et al. 2020). To look for such regions, we used a local PCA approach (Li and Ralph 2018) and identified a total of 47 outliers (**Supplementary Figures 10, 11**), ranging in size between 550kb and 25.7Mb (mean 3.9Mb) and covering in total 21.4% of the genome. Importantly, these regions contain 31.6% of *Δ*(*r*) outliers, and thus contribute disproportionately to the observed differences in recombination rates between the ecotypes (**Table 1**). Consistent with this, we found that the signal of recombination rate evolution measured by PRDI is about 10% stronger in the local PCA outliers than in the rest of the genome (**Supplementary Figure 12**). At the same time, it should be emphasized that there is also a strong PRDI signal of recombination rate evolution outside of these blocks (**Supplementary Figure 12**).

Large haplotype blocks can be the result of lack of recombination between alternative haplotypes segregating in Lake Masoko. We hypothesized that, on average, recombination would be reduced in the ecotype with the higher proportion of individuals who are heterozygous for such non-recombining haplotypes. Therefore, for each local PCA outlier region, we clustered the individuals based on individual heterozygosity, that is, the proportion of heterozygous sites per individual (H_ind_). An example of local PCA outlier region associated with differences in recombination is shown in **Figure 4**, with strong benthic vs. littoral clustering by H_ind_ illustrated in **Figure 4C**.

**Figure 4:**
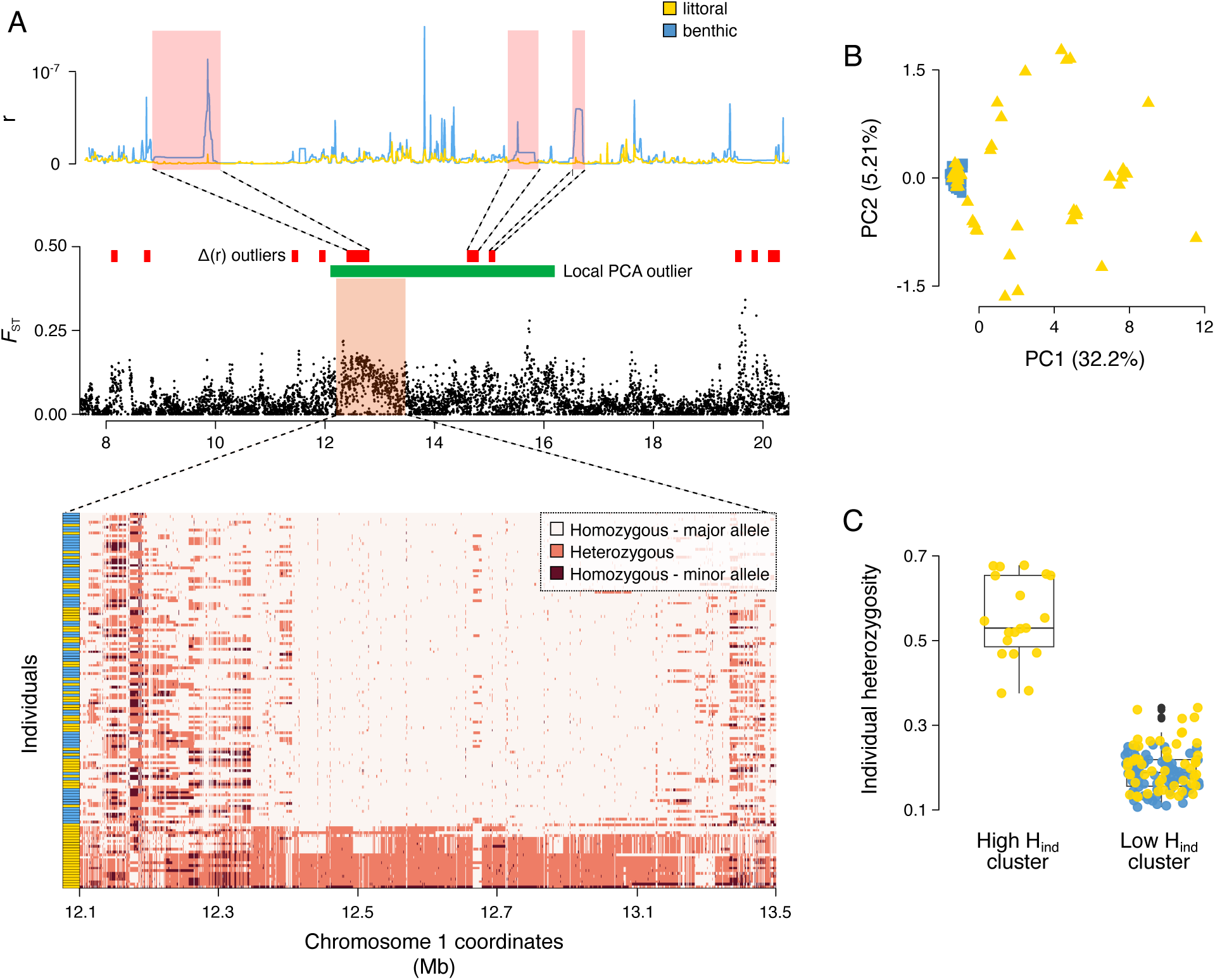
A local PCA outlier associated with difference in recombination. (**A**) An example region of chromosome 1 (LS420019.2) illustrating the link between a large haplotype block, *F*ST, and recombination rate evolution. Top: average recombination rates in the two ecotypes. There is lower recombination in the littoral ecotype in this region. Middle: *F*ST is moderately elevated in the haplotype block region. Bottom: Genotypes at SNPs with minor allele frequency > 5%. (**B**) Local PCA in the haplotype block region from (A). The genetic structure within this region differs considerably from the PCA obtained from whole-genome data (Figure 1B). (**C**) Clustering based on individual heterozygosity (Hind) - i.e. the proportion of variable sites that are heterozygous in an individual – in the haplotype block region from (A). All the individuals in the cluster with high Hind belong to the littoral ecotype.

As predicted, across the 47 haplotype blocks we found significant positive association between Δ(*r*), the net recombination distance between ecotypes, and the degree of ecotype clustering by H_ind_ (**Figure 5A**). Additionally, PRDI was substantially elevated in the haplotype blocks with significant of ecotype clustering by H_ind_ (**Supplementary Figure 12**). In contrast, the association of Δ(*r*) with allele frequency divergence (mean *F*_ST_ per haplotype block) with was less pronounced and not statistically significant (**Figure 5B**).

**Figure 5:**
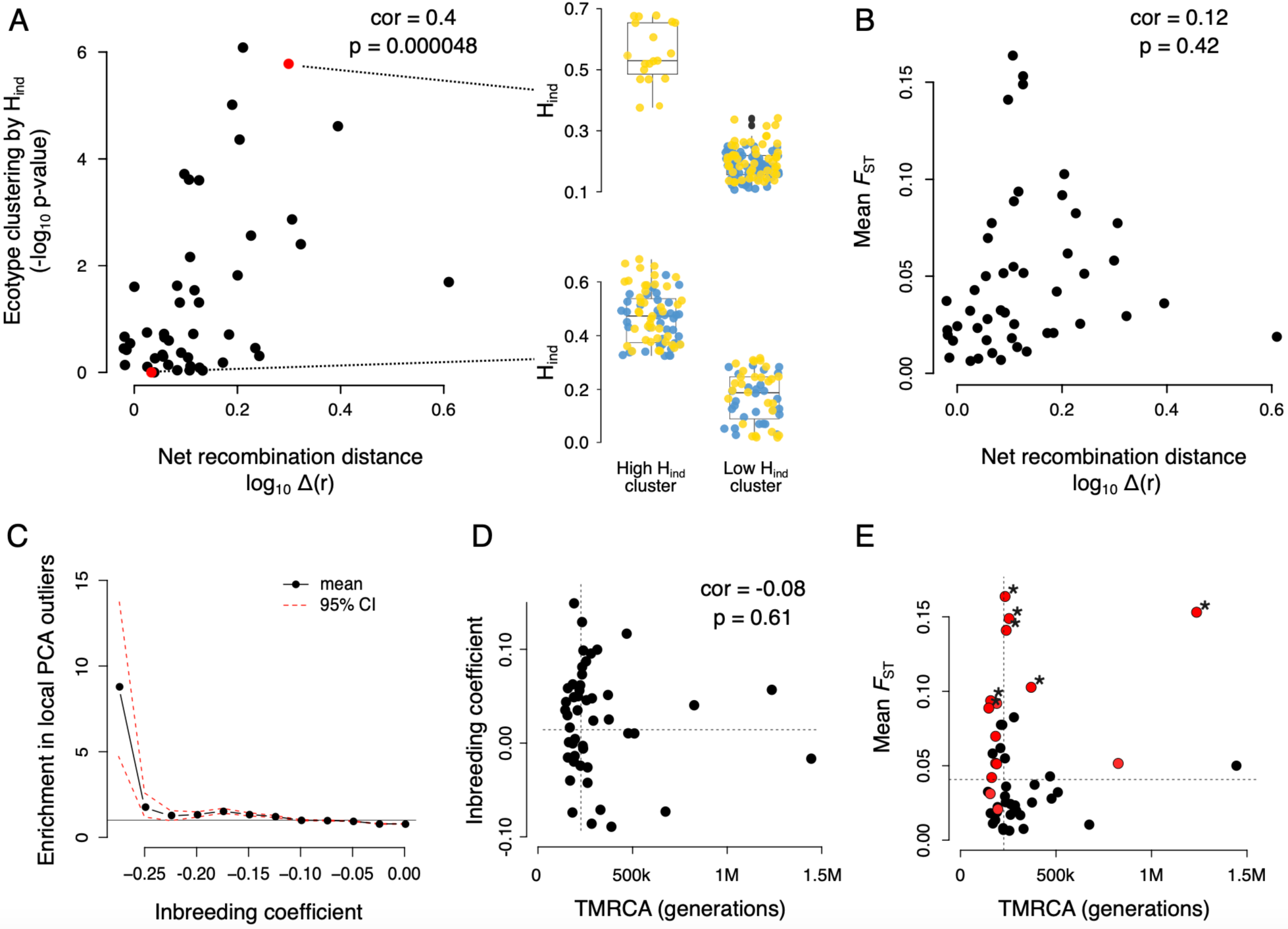
Characterizing haplotype blocks and their link with recombination in Lake Masoko. (A) Relationship, across the 47 local PCA outliers, between the degree of ecotype clustering by Hind and net between-ecotype recombination map distances. We highlighted examples of a strong (top) and weak (bottom) ecotype clustering by Hind. (B) Relationship between mean *F*ST and net between-ecotype recombination map distances. (C) For each value of inbreeding coefficient, we show the relative enrichment of SNPs in the PCA outlier regions in comparison with the control regions. (D) Relationship between inbreeding coefficient and TMRCA. The horizontal dashed line represents the mean inbreeding coefficient calculated for the control regions while the vertical dashed line represents an estimate of genome wide average TMRCA. (E) Relationship between TMRCA and mean *F*ST. The local PCA outliers that contain island(s) of differentiation are colored in red. We used stars to mark the regions with significantly elevated FST (permutation test p-value < 0.05).

Next, we focused on gathering evidence regarding the nature and origin of individual haplotype blocks. Local PCA outliers could be explained for example by linked selection, recent admixture from outside of Lake Masoko, or by locally low recombination rates. We found that some of the regions we identified by local PCA show signatures characteristic of polymorphic inversions, including (i) long haplotypes with consistent sharp edges in multiple individuals, (ii) distinct homozygote vs. heterozygote clusters in PCA, and (iii) unusually high values of individual heterozygosity (**Supplementary Figures 13, 14**).

To investigate if the heterozygous state is overrepresented within local PCA outliers, which would be consistent with a form of balancing selection, we compared the distribution of inbreeding coefficient (F) per SNP against SNPs from a set of control regions (**Methods**). We found a significant enrichment of SNPs with negative *F* – that is, excess of heterozygotes – in the local PCA outliers, with up to 9-fold enrichment for the SNPs at the lowest values of *F* (**Figure 5C**). We expected that the haplotype blocks with the lower value of inbreeding coefficient may have been maintained as polymorphism for a long time through the effect of balancing selection. Therefore, for each PCA outlier we estimated the time to the most recent common ancestor (TMRCA). Perhaps surprisingly, we found that the ancestry of most of these regions was of a similar age as the genome wide average, with only a few that were clearly much older (**Figure 5D**). There was a weak and not statistically significant trend of older regions having a lower inbreeding coefficient. Therefore, we conclude that the effect of balancing selection on maintaining polymorphic haploblocks appears to be limited in the Lake Masoko ancestry.

Finally, we investigated in more detail the link between allele frequency divergence and the local PCA outliers in the Lake Masoko. We found that only seven out of the 47 regions had a significantly elevated average level of *F*_ST_; of these, there were six regions of average age and one old region whose ancestry dates back almost 1.5M generations ago (**Figure 5E**). Islands of differentiation – also referred to as highly diverged regions (HDRs), defined as in ref. (Malinsky et al. 2015) – also did not appear in many of the local PCA outliers. While four of the local PCA outliers contained more than ten HDRs each, in total only 15 local PCA outliers contained at least one HDR (**Figure 5E**; **Supplementary Table 2**).

## Discussion

The landscape of recombination across the genome is not static but evolves through time. In this study, we undertook a holistic investigation of recombination rate evolution between two ecotypes that diverged very recently, in sympatry with gene flow, and adapted rapidly across multiple phenotypic traits to new lake environments (Malinsky et al. 2015). Given ecotype divergence at many – at least a hundred or so – genomic loci (Malinsky et al. 2015), we could expect that recombination, or the lack thereof, would be important in bringing together and keeping together the alleles that are beneficial in each environment (Ortiz-Barrientos et al. 2016; Todesco et al. 2020; Schluter and Rieseberg 2022; Battlay et al. 2023). Our findings reveal, characterize, and quantify substantial differences in population recombination rates between the ecotypes, complementing previous studies of divergence in ecology, mate choice, and allele frequencies (Malinsky et al. 2015), methylation (Vernaz et al. 2022), gene expression (Carruthers et al. 2022), and in sex determination (Munby et al. 2021).

The use of LD-based maps, integrating over a large number of recombination events throughout the ancestry of the ecotypes, allowed us to analyze recombination landscape evolution at a fine scale. Despite the challenges inherent in such comparisons, we used subsampling (replicate maps from the same ecotype) and extensive simulations to derive meaningful measures of evolution in population landscapes. We show how the recombination differences are distributed across the genome, how they are linked to allele frequency divergence (*F*_ST;_ *Δ*(*π*)) and to large haplotype blocks identified by local PCA.

As we expected, we found a link between recombination rate divergence and regions of high allele frequency divergence. This link can be expected for a variety of reasons – for example, local recombination rates can impact inference of *F*_ST_ (Booker et al. 2020), they partly determine the extent of background selection which can affect *F*_ST_ (Burri et al. 2015; Matthey-Doret and Whitlock 2019), and they play a major role in in the formation of genomic islands of divergence during speciation with gene-flow (Feder et al. 2012; Cruickshank and Hahn 2014). Moreover, selection on genotypes (e.g., incomplete selective sweeps) is directly reflected in LD-based recombination maps (in contrast to gamete-based or pedigree-based maps) (Coop and Przeworski 2007; Peñalba and Wolf 2020). Given these considerations, it is surprising that allele frequency divergence does not appear to be the main driver of the observed recombination rate differences. For example, only 13.7% of significant differences in recombination, i.e. *Δ*(*r*) outliers, are co-located with the top 10% of *F*_ST_ (**Table 1**), and PRDI is substantially positive in regions where *F*_ST_ and *Δ*(*π*) are zero.

A stronger link was identified between recombination rate evolution and local PCA outliers, which are characterized by large haplotype blocks and comprise more than a fifth of the genome. Some of these blocks carry clear signatures characteristic of inversions. However, it remains to be seen what mechanisms give rise to the remaining blocks where the haplotype structure and local relationships among individuals are more complex. In many cases we see many clusters in the local PCA, which is consistent with signatures of multiple overlapping inversions segregating locally in the population (Faria et al. 2019a). There is growing evidence that haplotype blocks caused by large inversions, are present in many species (Wellenreuther and Bernatchez 2018), often comprise considerable proportions of the genome, and have clear links to adaptation and diversification in both animals (Faria et al. 2019a; Harringmeyer and Hoekstra 2022; Reeve et al. 2023; Blumer et al. 2024) and plants (Todesco et al. 2020; Battlay et al. 2023). Therefore, future work should focus on exploring the dynamics of inversions in the Lake Masoko system in greater detail.

We have focused on large haplotype blocks because the local PCA approach facilitates their study from short read data in a population genomic context. However, the recombination suppression effect of structural variants does not depend on the size of the variant region, and other types of structural variation can also suppress recombination (Kent et al. 2017; Rowan et al. 2019; Mérot et al. 2020). Therefore, it is likely that shorter structural variants are responsible for at least some of the remaining *Δ*(*r*) outliers which are not accounted for in our current study. Further reduction in the cost of long-read sequencing will, among other benefits, enable more unbiased population-scale analyses of structural variants and their roles in evolution of recombination landscapes (Coster et al. 2021).

We found substantial recombination landscape evolution where the ecotypes cluster by individual heterozygosity levels (**Figure 5A**, **Supplementary Figure 12**). This could be because inversions prevent crossover formation only in the gametes of heterozygous individuals (Faria et al. 2019b). A recent study using pedigree-based recombination maps raises an additional possibility. Venu et al. (2024) found that regions with higher heterozygosity have lower recombination rates due to haplotype incompatibilities between diverging ecotypes. Therefore, the same mechanism could play a role in the divergence of recombination landscapes seen in Lake Masoko.

More general and important open questions concern the nature of selection on recombination. Which of the observed differences in population recombination rates are a result of changes in the distribution of crossovers during gamete formation? And which changes are an indirect effect of subsequent selection for or against specific genetic variants and recombinant haplotypes? These questions are not possible to answer conclusively with LD-based estimates alone. A future comparison of our LD-based maps against recombination landscapes obtained by sequencing of gametes and / or individuals related by pedigrees will shed further light on this question (Peñalba and Wolf 2020).

Our finding of rapid recombination rate evolution, while consistent with some previous studies (Shanfelter et al. 2019; Déserts et al. 2021), seems to be in conflict with the current paradigm of evolutionary stability of recombination landscapes in species lacking the PRDM9 mechanism (Lam and Keeney 2015; Singhal et al. 2015). However, this is not as surprising as it may appear because it is common that different tempos of molecular evolution are observed between micro- and macro-evolutionary timescales (Rolland et al. 2023). For example, recombination suppression by inversions may be a temporary phenomenon and may disappear once one of the inversion alleles rises to fixation. At the same time, evidence is emerging that even in many species with an intact PRDM9 mechanism, a large fraction of recombination can take place outside of PRDM9 directed hotspots, so the dichotomy of mechanisms may not be as clear as previously thought (Hoge et al. 2023; Joseph et al. 2023).

Adaptation and organismal diversification are increasingly seen as multidimensional and combinatorial, typically with involvement of multiple polygenic traits and epistasis (Marques et al. 2019; Barton 2022; Yeaman 2022), and the relative genetic distances between the loci involved constitute key parameters. Comparative studies are starting to shed light on recombination landscape evolution across populations and species with different demographic histories, genomic architectures, ecological contexts, and divergence times. However, this is still typically done at a rough Megabase-scale resolution (Haenel et al. 2018; Brazier and Glémin 2022). We adopted the LD-based approach, enabling us to infer fine-scale rates and show that they can evolve rapidly. We envisage that the large and growing amount of population genomic data available will enable construction and comparisons of many LD-based maps, such as in our current study. Together with advances in gamete typing and pedigree-based methods, this will make recombination rates and their fine-scale evolution into integral parts of future genomic studies of adaptation and speciation.

---

**Supplementary Figure 1:**
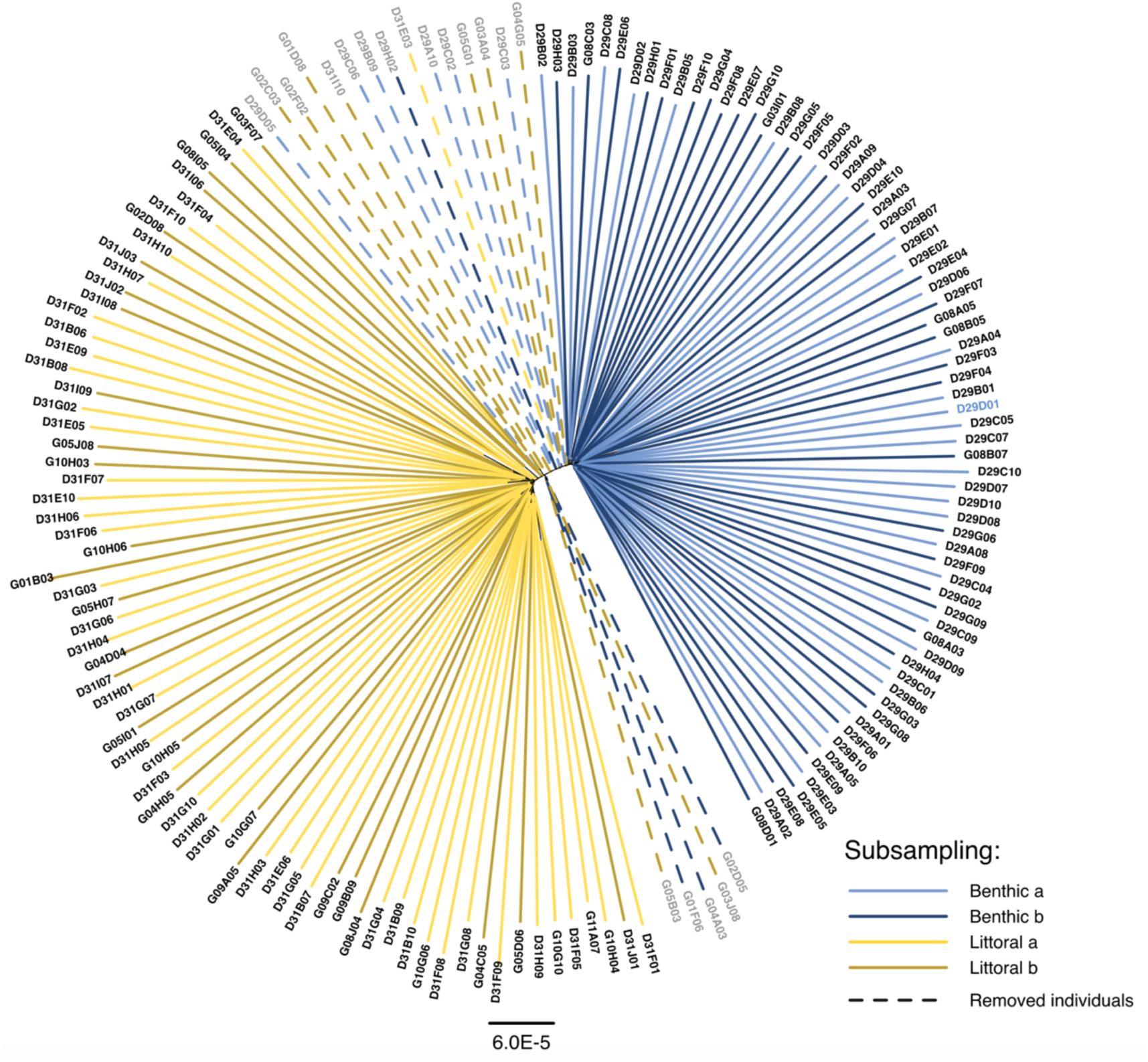
Neighbor joining tree based on the number of differences in polymorphic sites between individuals. Each tip represents one individual. We used this tree and the PCA showed in Figure 1 to determine which intermediate individuals to discard from downstream analyses. After discarding the intermediate individuals (indicated by dashed lines), the remaining samples constitute two monophyletic groups in this tree.

**Supplementary Figure 2:**
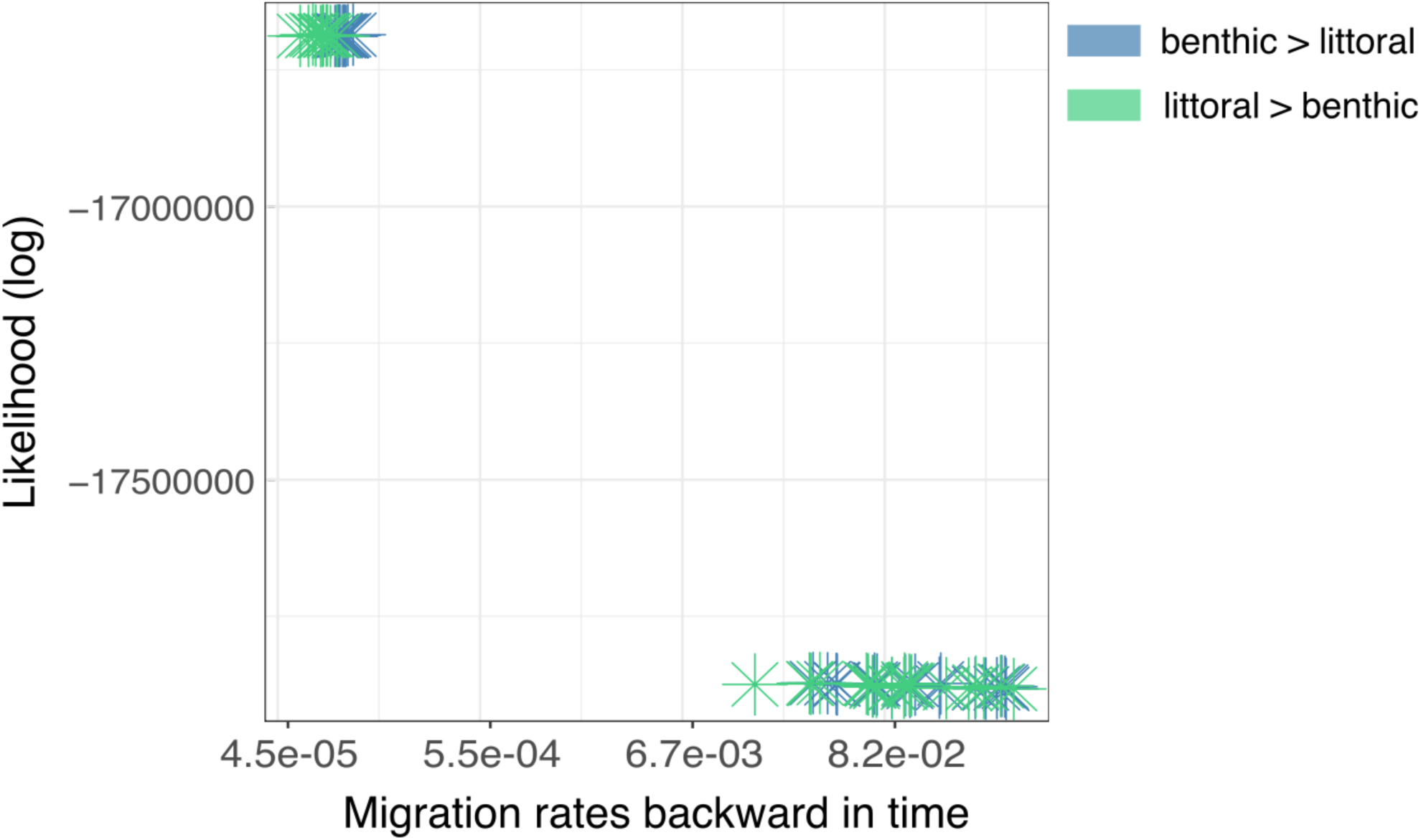
Estimating migration rates with fastsimcoal2. Each point represents an estimate from an independent run with different starting parameters. The estimates with lower migration rates (top left corner) have superior likelihoods to the estimates in the bottom right. In the main text, we report the overall maximum likelihood estimates. However, we note that given the number of intermediate individuals in our sample (presumably recent hybrids), the maximum likelihood rates are lower that we would expect and the results with the lower likelihood should not be completely discounted.

**Supplementary Figure 3:**
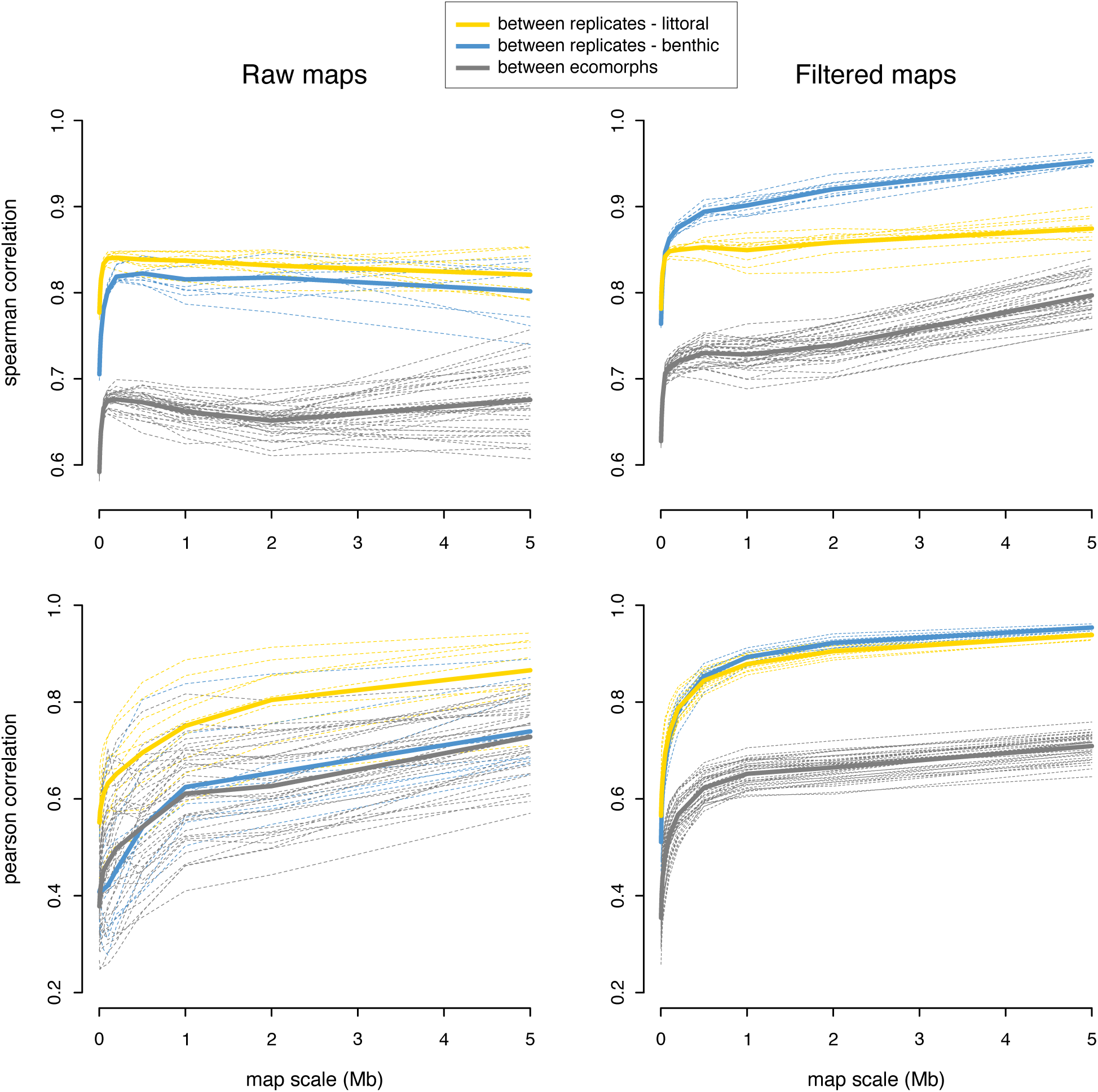
Correlation between recombination maps at different scales. We see that correlations in the between ecotype comparisons are considerably lower than correlations for the within-ecotype replicates. This holds across all genomic scales (2kb to 5Mb) for Spearman correlation, across almost all genomic scales for Pearson correlation, and across bootstrap replicates of genetic maps obtained by randomly sub-sampling individuals for each ecotype (dashed lines). We also see that the filtered maps (after masking 30.6% of the genome; see Materials and Methods) show considerably less inference noise than the raw maps, making the results even clearer.

**Supplementary Figure 4:**
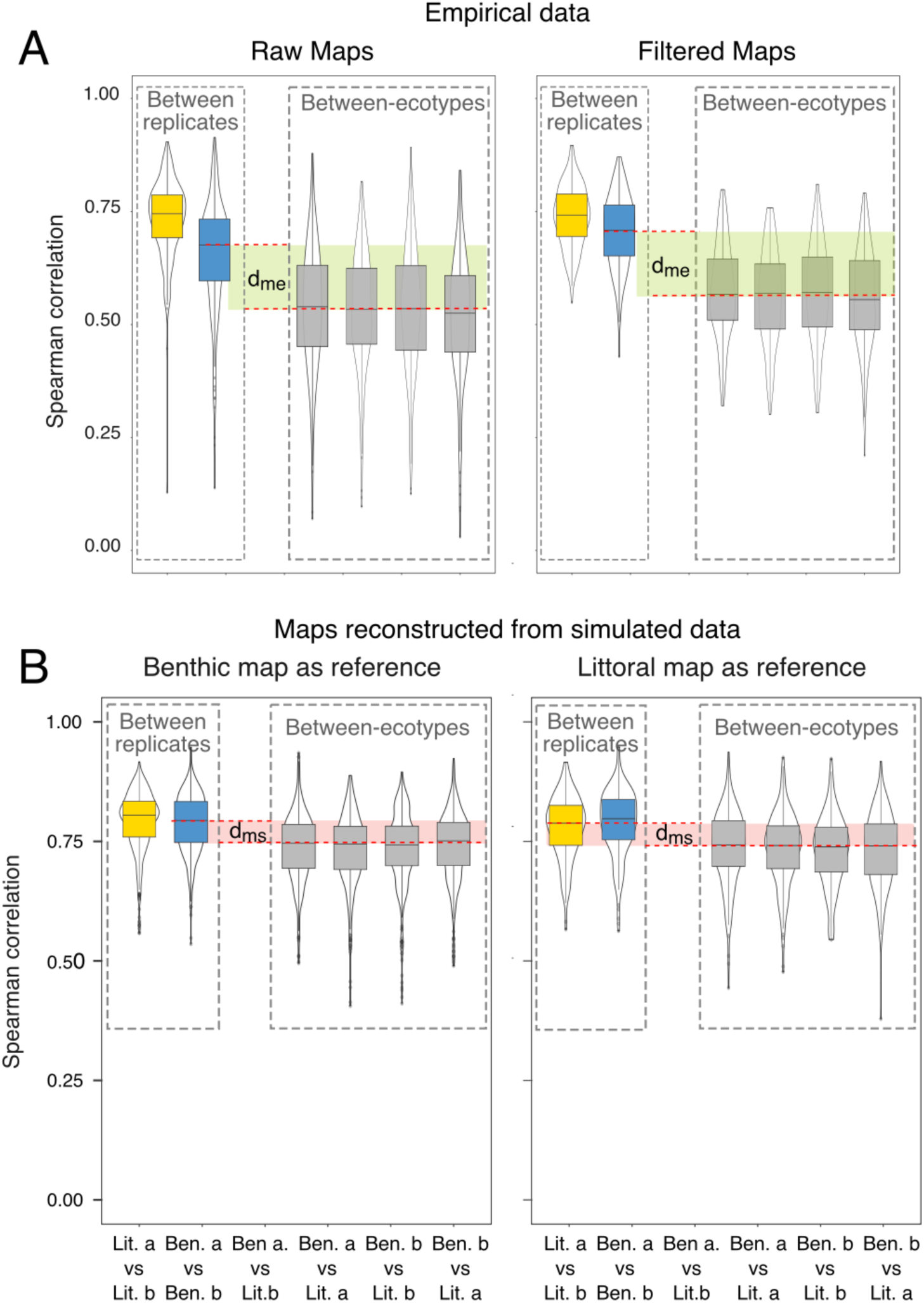
Confirming the robustness of the PRDI estimate of recombination landscape evolution. **(A)** A comparison of recombination map correlations derived from the raw maps (equivalent to Figure 2A) and maps that have been carefully filtered to exclude regions where recombination inference might have been error prone (Filtered Maps; see main text for details). Importantly, the d_me_ estimate, which is the first term in the PRDI calculation, is virtually identical. **(B)** A comparison of recombination map correlations for maps derived from simulated data. Regardless of which empirical map was used as an input/reference for the simulations, we got virtually identical estimates of d_ms_, the second term in the PRDI calculations.

**Supplementary Figure 5:**
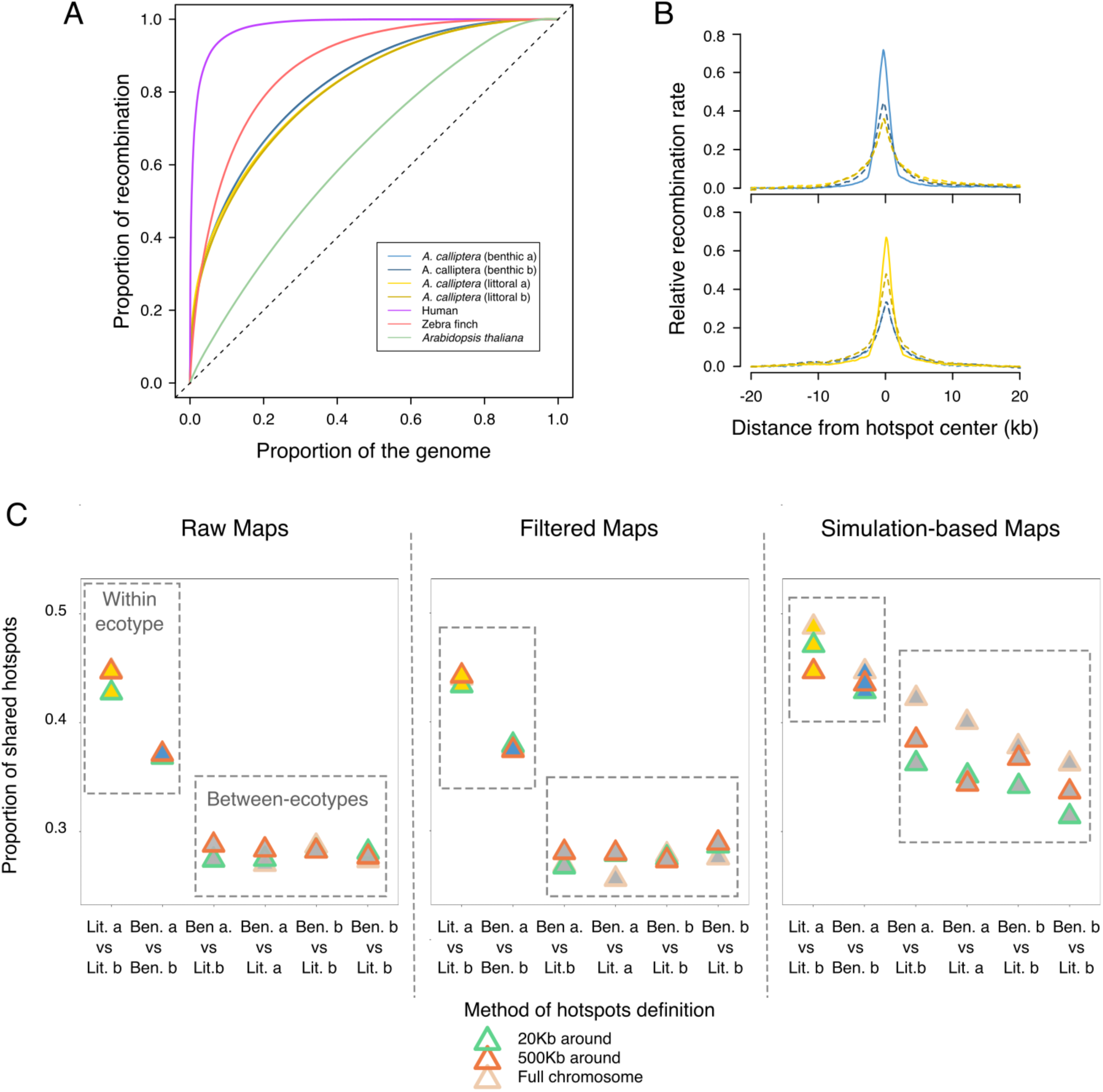
Characterizing recombination hotspots and their evolution in Lake Masoko. Blue and yellow colors refer to the benthic and littoral ecotypes respectively. **(A)** Empirical cumulative distribution function of for recombination fraction as a function of proportion of genome covered. **(B)** Scaled recombination rates around hotspots defined in the benthic a (top) and littoral a (bottom) subsamples (solid lines) and the scaled rates at these position in the other subsamples (dashed lines). Hotspots in (B) are defined against a 1Mb background. The light color (blue and yellow) corresponds to the subsamples a (in benthic and littoral) and the dark color to the subsamples b. **(C)** The proportion of overlapping hotspots between recombination maps, based on three different background sizes - these three methods differ on the size of the genomic background used as baseline (e.g. 20kb, 500kb and full chromosome). We observe very similar proportion of overlapping hotspots with different the different

**Supplementary Figure 6:**
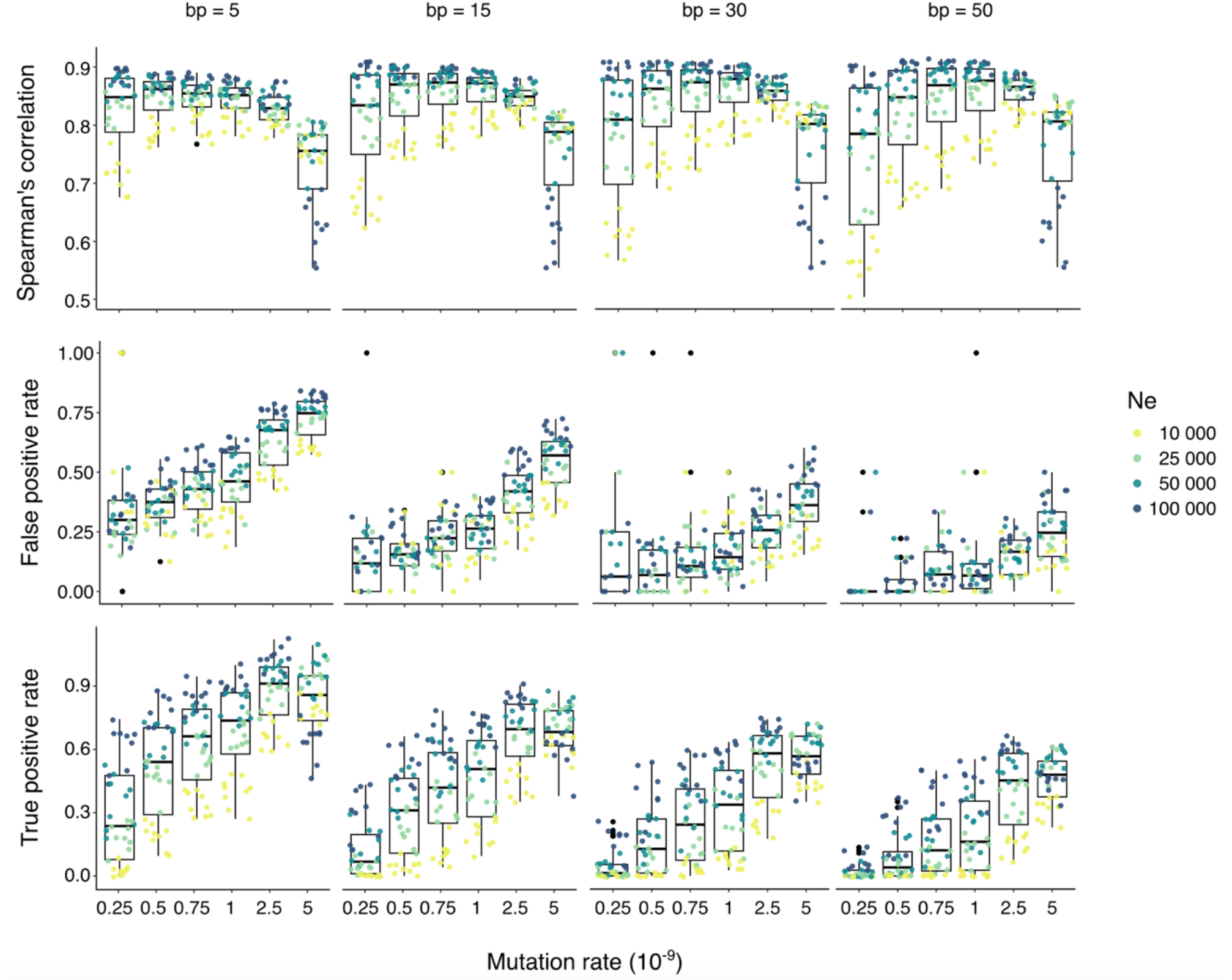
Inference of recombination maps and hotspots from coalescent simulations. We used `msprime` to simulate ancestral history with mutations based on different mutation rates (*μ*) and effective population sizes (*N_e_*). We inferred for each VCF the recombination landscape with *pyrho* using different block penalty values (bp = 5 to bp = 50). Here, we plot (i) the Spearman correlation, (ii) the false positive rate in hotspot discovery and (iii) the true positive rate in hotspot discovery between the recombination maps inferred from `msprime` simulation and the reference map. Recombination inference clearly depends on genetic diversity *π* = 4*N*_*e*_*μ*, and performs generally better when the level of *π* is higher.

**Supplementary Figure 7:**
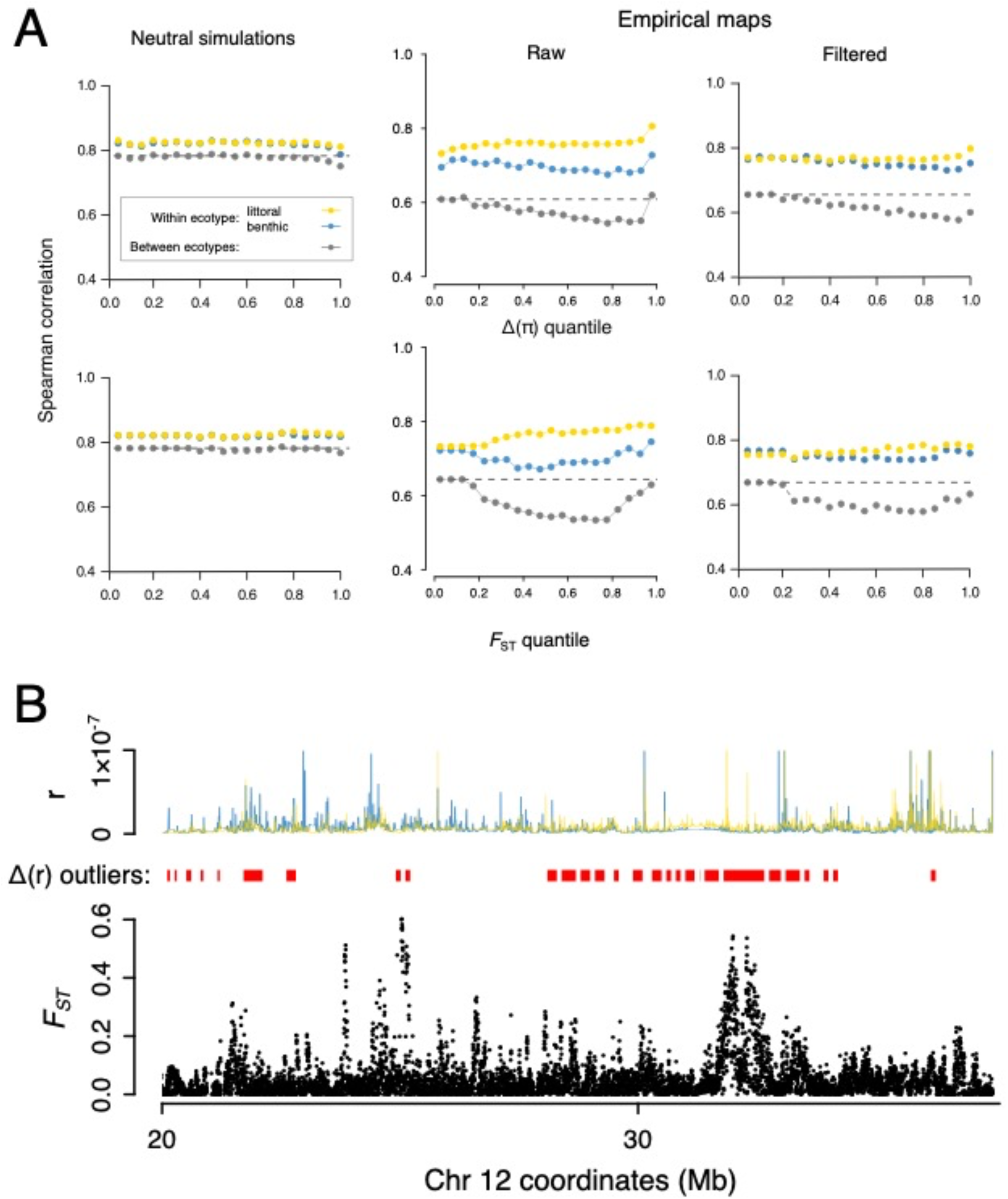
The relationship between allele frequency divergence and recombination divergence. (**A**) Spearman correlation measured between recombination maps as a function of benthic-littoral *F*_ST_ and of Δ(π). There is no relationship in neutral coalescent simulations. Empirical maps, both raw and filtered, show a consistent result of always lower-than-expected correlations between ecotypes and a consistent trend of lower benthic vs. littoral correlations in regions of greater *F*_ST_ and of Δ(π). (**B**) An illustrative example. While some Δ(r) outliers – regions of significantly divergent recombination – co-locate clearly with elevated *F*_ST_ others do not and there are also clear *F*_ST_ peaks in regions where benthic and littoral recombination rates appear to be similar.

**Supplementary Figure 8:**
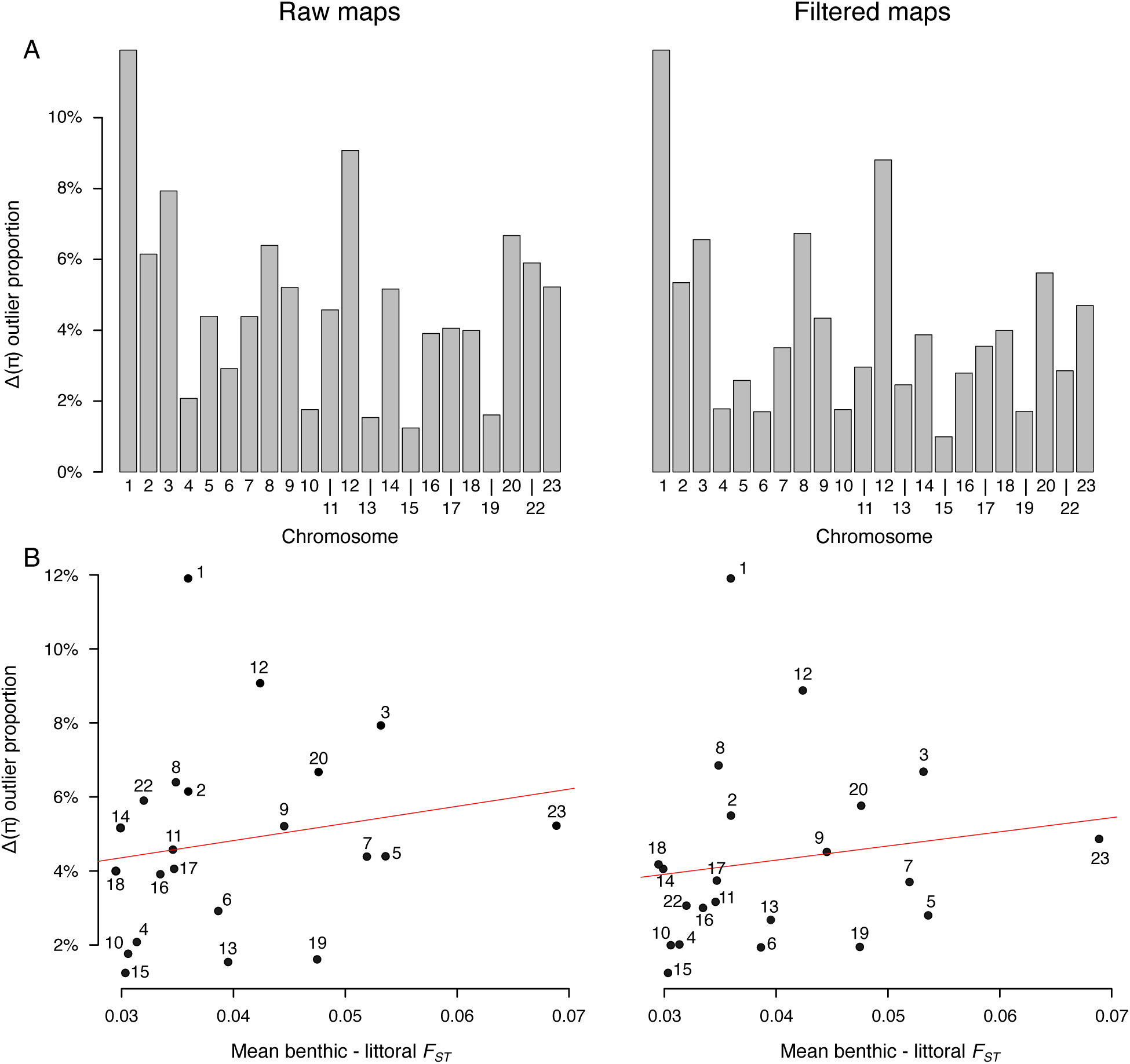
The distribution of *Δ*(*r*) outliers across chromosomes. (**A**) The proportion of each chromosome assigned as *Δ*(*r*) outliers, i.e., regions with rapidly changing recombination landscapes. (**B**) The relationship between *Δ*(*r*) outlier proportion and mean *F_ST_* per chromosome.

**Supplementary Figure 9:**
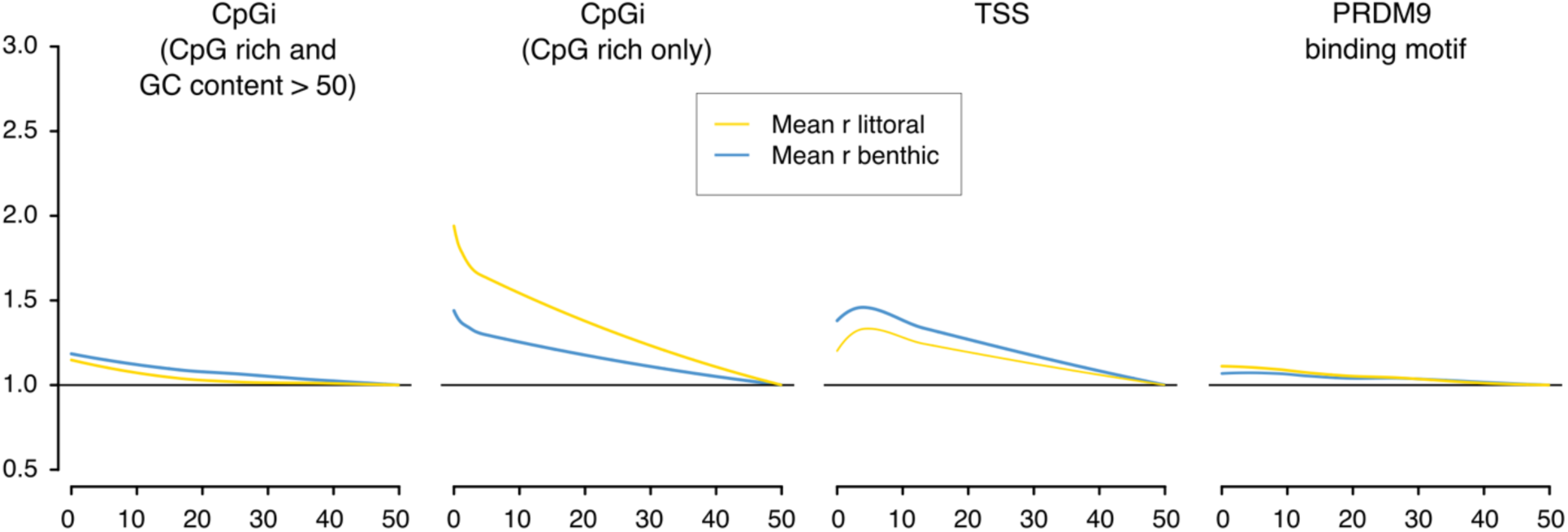
Recombination landscapes in Lake Masoko have PRDM9-independent profiles. Relative recombination rates in 2kb windows are higher near CpG islands (GpGi) and transcription start sites (TSS) but are independent of the binding motif associated with the incomplete PRDM9 ortholog presents in this species. <u>CpG islands were defined with and without condition of</u> <u>50% minimal GC content (see Materials and Methods)</u>.

**Supplementary Figure 10:**
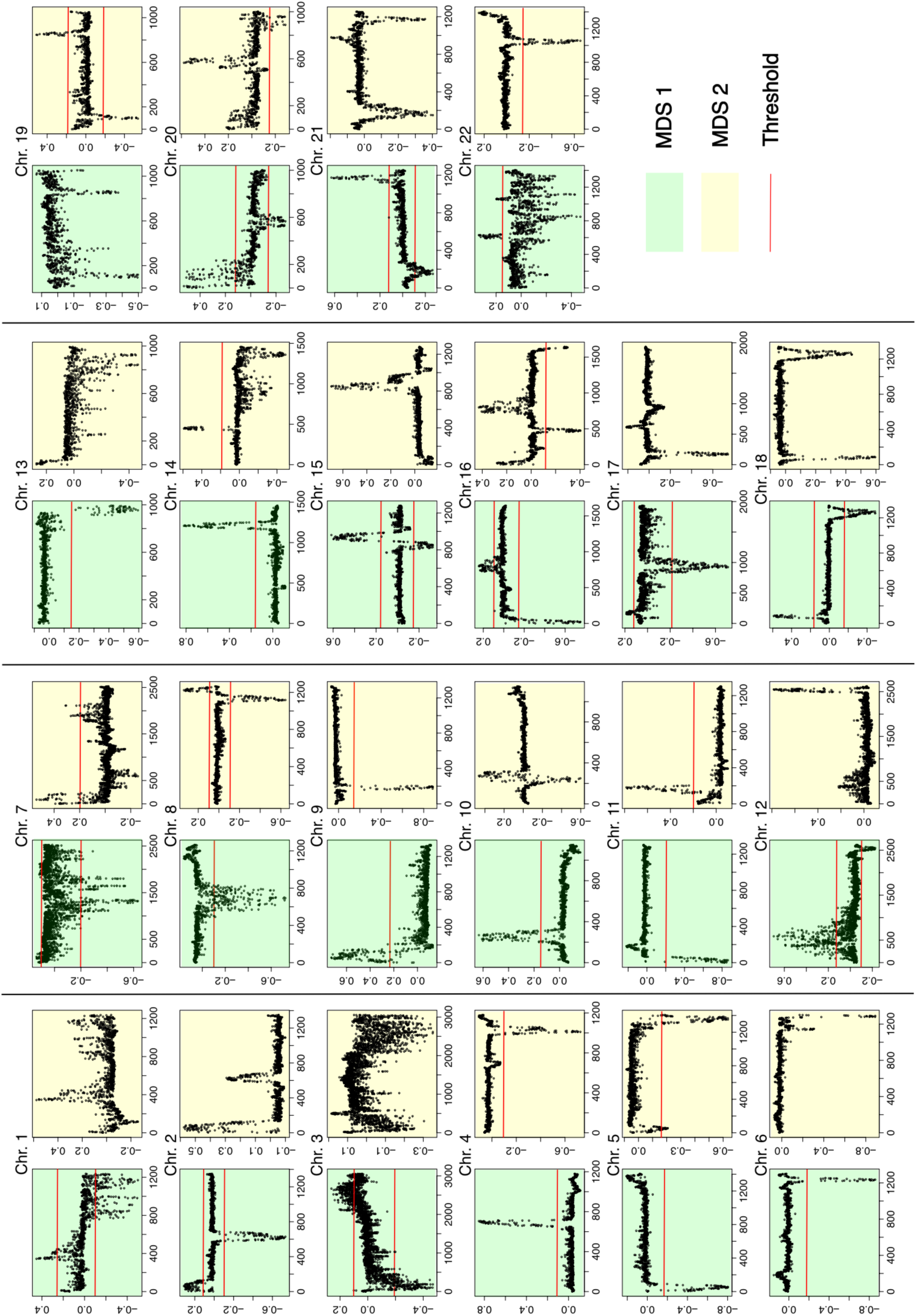
Detection of genomic regions showing clearly distinct population structure. Each datapoint corresponds to a local PCA for a 100 SNP genomic window. The (dis-)similarity between population structure shown in different local PCA windows is measured by multidimensional scaling (MDS): the greater the MDS deviation from the chromosome average, the more distinct the local population structure. We visually inspected two MDS coordinates (MDS 1 and MDS 2) to select thresholds for each chromosome, identifying 47 distinct regions.

**Supplementary Figure 11:**
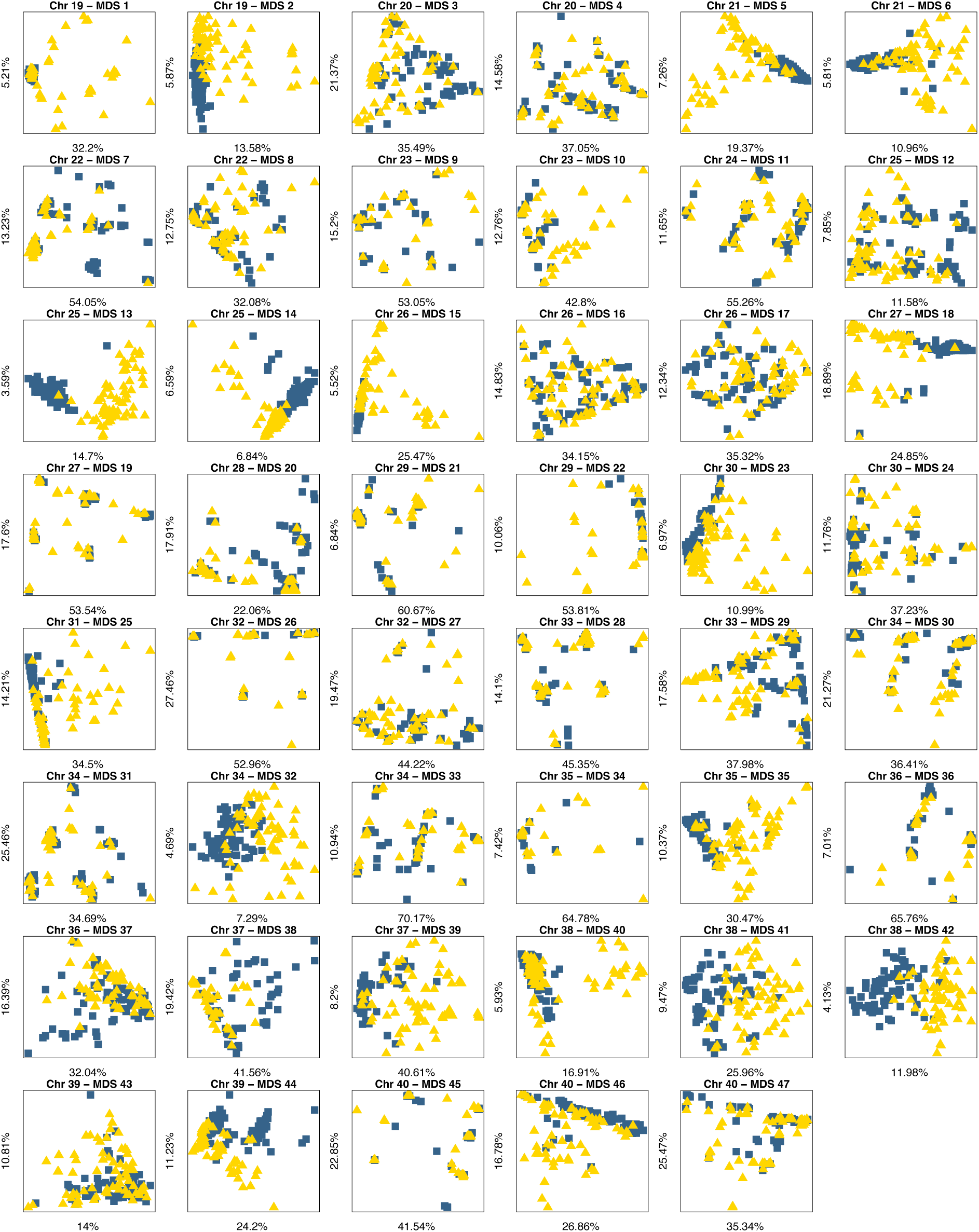
Local PCAs within the 47 haplotype blocks identified with lostruct. Each blue square represents a benthic individual, and each yellow triangle a littoral individual. The patterns of population structure are very diverse, with some showing clear differences between the ecotypes. In all cases, the first two principal components explain substantially greater proportion of total genetic variation than in the whole genome PCA shown in Fig 1B.

**Supplementary Figure 12:**
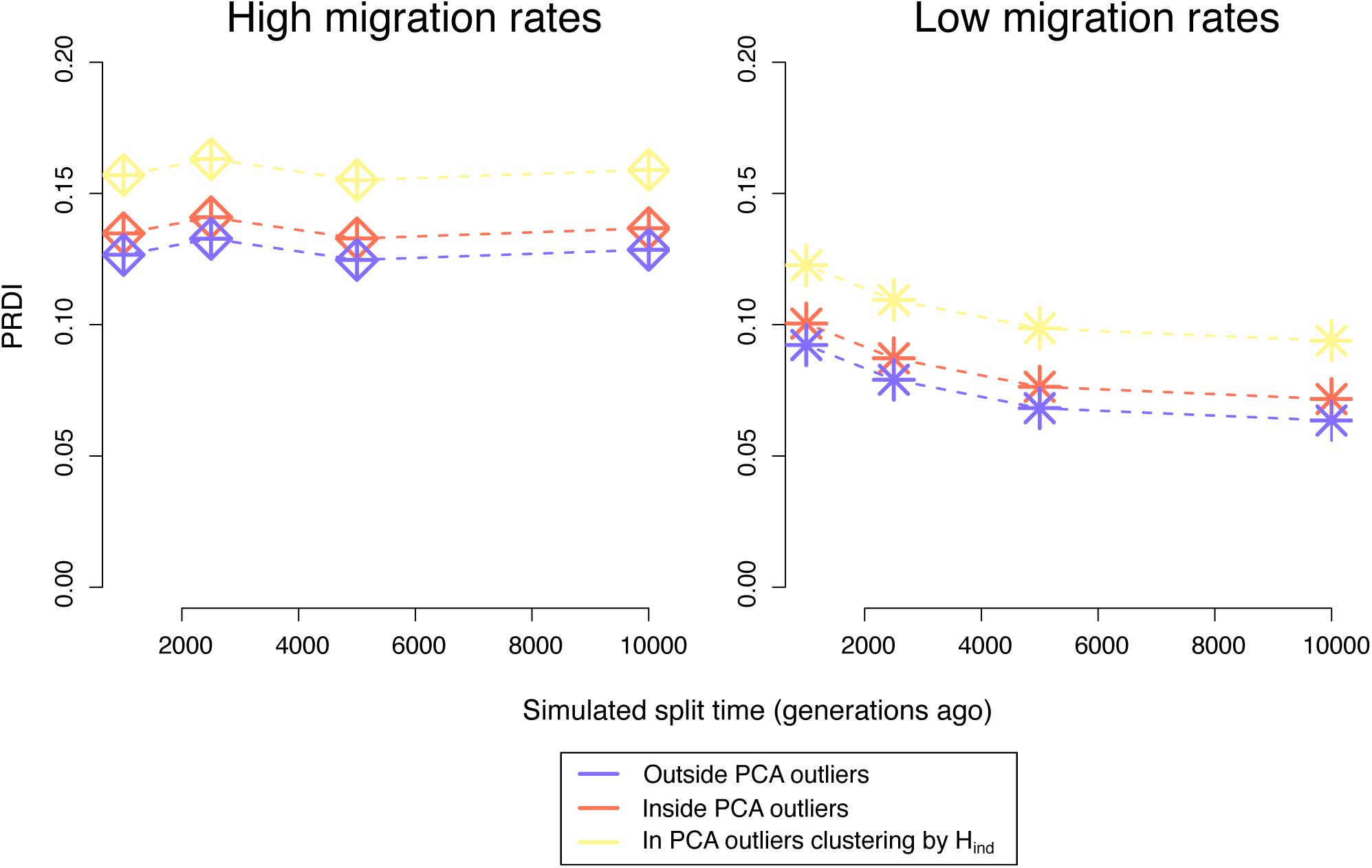
The relationship between ecotype divergence in recombination and local PCA outliers. The Population Recombination Divergence Index (PRDI) is about 10% higher in local PCA outliers than outside of these blocks. PRDI is further elevated in the local PCA outliers where the ecotypes cluster based on individual heterozygosity (H_ind_). The same conclusions are obtained regardless of the split times and/or migration rates used in the neutral simulations that form a part of PRDI estimation.

**Supplementary Figure 13:**
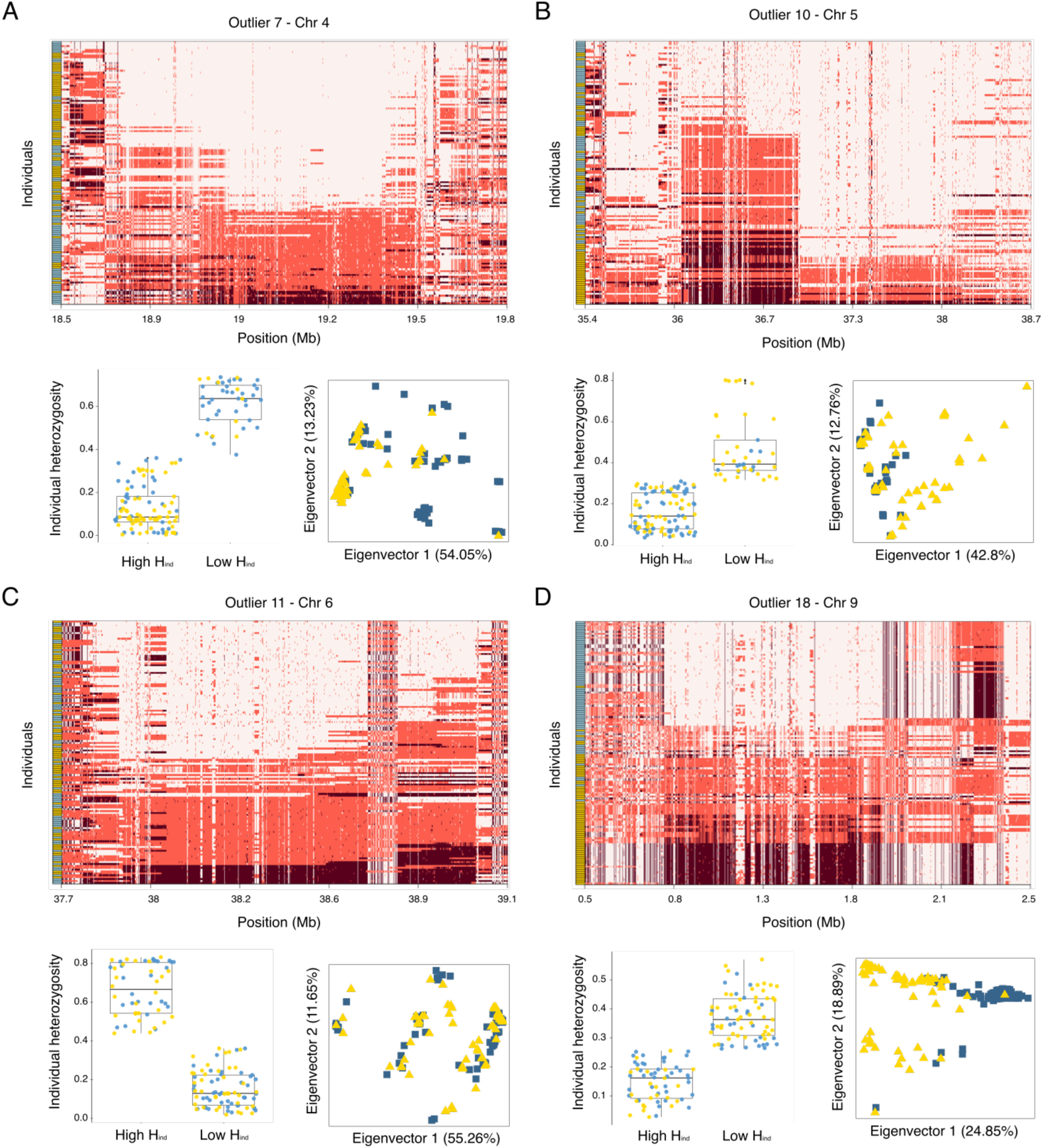
Examples of large haplotype blocks with typical inversion signatures. For four `lostruct` outliers we show individual genotypes (with minor allele frequency > 5%; color scheme as in **Figure 4A**), results of k-means clustering (K=2) of individuals based on their heterozygosity, and the local PCA for that region. We see sharp edges of haplotype blocks, individuals who are consistently heterozygous for long stretches, and individuals falling clear clusters in the PCA, all characteristic signatures of inversion. However, each of these `lostruct` outliers contains several different haplotype blocks. A more detailed study of history and mechanisms behind these haplotype blocks will require careful delineation of the boundaries of each of these haplotypes.

**Supplementary Figure 14:**
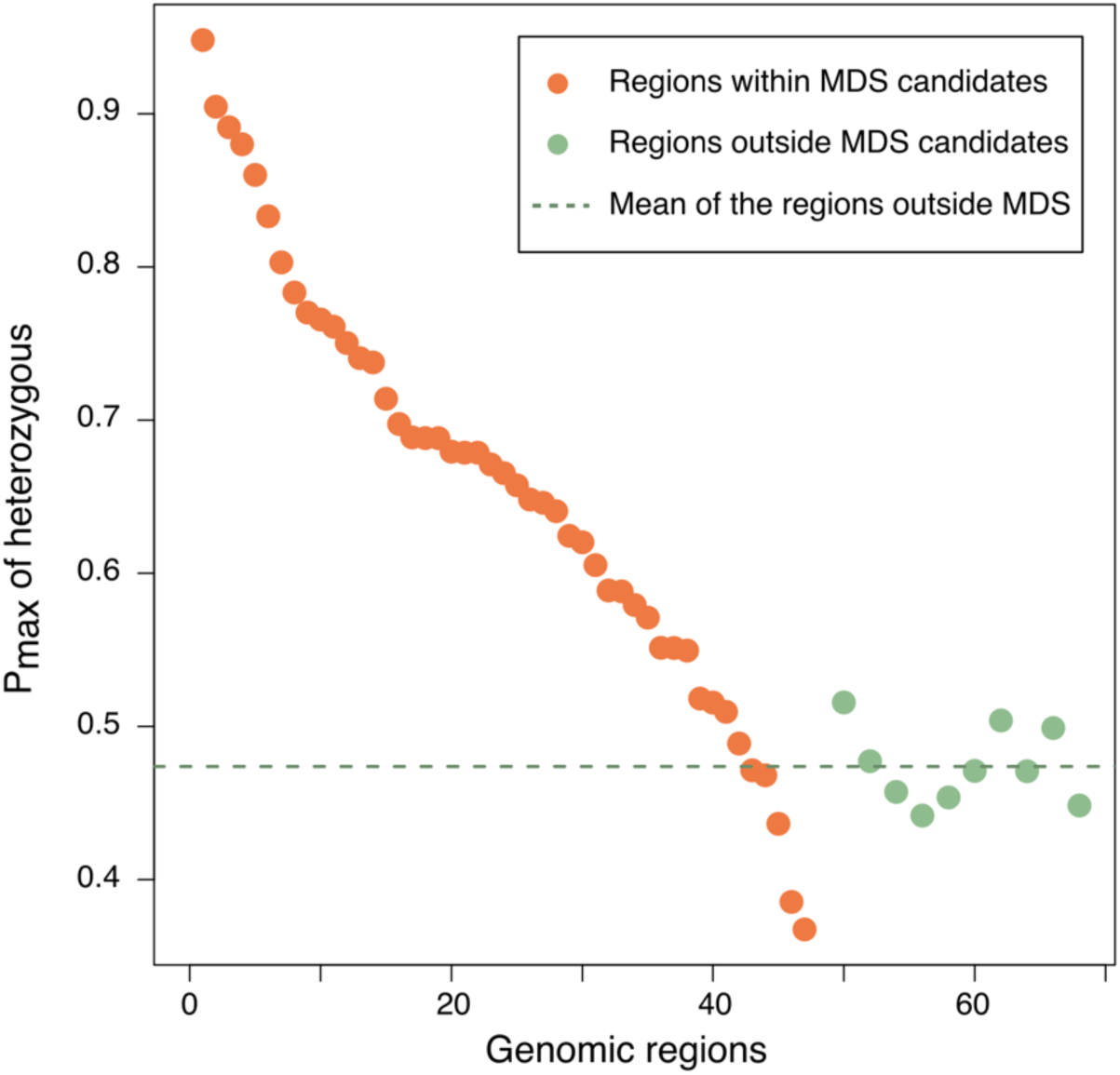
High levels of individual heterozygosity in regions identified as distinct by local PCA. We found that for 90% of the local PCA outlier regions, the maximum of individual heterozygosity (H_ind_) was higher than the mean of the equivalent measure in length-matched control regions.

**Supplementary Table 1:** The number of hotspot defined relative to the 3 different background that we used. . We defined a hotspot of recombination in a given genomic region if the recombination rate was at least 5 times higher than the recombination rate in the background region. The genomic area longer than 5kb were not considered as hotspots.

|  | *Number of hotspots* |  |  |
| --- | --- | --- | --- |
|  | Define on 20kb<br>around | Define on 500kb<br>around | Define on the<br>chromosome mean |
| **Benthic a** | 2350 | 2275 | 2132 |
| **Benthic b** | 2488 | 2341 | 2249 |
| **Littoral a** | 2086 | 2345 | 2378 |
| **Littoral b** | 2086 | 2328 | 2433 |

**Supplementary Table 2:** The genomic coordinates and key statistics for the 47 regions identified as local PCA outliers. The “cluster p-value” column refers to the ecotype clustering -log_10_ p-value shown in Figure 5A. HDRs refers to the number of highly diverged regions (HDRs; defined as in *(Malinsky et al. 2015)*) that overlap the local PCA outlier.

| chr | start | end | $F$ | $F_{ST}$ | $H_{ind}$ | cluster<br>p-value | $\log_{10} \Delta(r)$ | TMRCA | HDRs |
| --- | --- | --- | --- | --- | --- | --- | --- | --- | --- |
| 1 | 12102312 | 16189411 | 0.0166 | 0.0582 | 0.6882 | 5.78 | 0.2992 | 168146 | 0 |
| 1 | 26036869 | 41119330 | 9e-04 | 0.0421 | 0.5495 | 5.01 | 0.1899 | 163307 | 1 |
| 2 | 19465 | 1831057 | 0.0732 | 0.055 | 0.6711 | 0 | 0.1073 | 232967 | 0 |
| 2 | 14469831 | 18569566 | -0.04 | 0.0112 | 0.6787 | 0.04 | 0.1326 | 169577 | 0 |
| 3 | 1119826 | 10621518 | 0.0513 | 0.1027 | 0.3854 | 4.36 | 0.2042 | 369977 | 15 |
| 3 | 37891964 | 63560498 | 0.0404 | 0.0515 | 0.3675 | 1.35 | 0.0882 | 825661 | 3 |
| 4 | 18591787 | 19609472 | 0.0568 | 0.1531 | 0.7405 | 3.6 | 0.1257 | 1235990 | 1 |
| 4 | 27040234 | 29900969 | -0.0241 | 0.0069 | 0.6787 | 0.08 | 0.0835 | 224469 | 0 |
| 5 | 61769 | 1324715 | 0.0616 | 0.008 | 0.8801 | 0.42 | -0.015 | 223426 | 0 |
| 5 | 35372227 | 38655485 | 0.0955 | 0.0825 | 0.803 | 2.57 | 0.2263 | 279811 | 0 |
| 6 | 37886749 | 39005213 | 0.0457 | 0.0063 | 0.8331 | 0.1 | 0.0254 | 255967 | 0 |
| 7 | 44298 | 5486542 | 0.0239 | 0.0197 | 0.5156 | 0.14 | -0.018 | 292390 | 0 |
| 7 | 23358419 | 45879790 | 0.0585 | 0.0937 | 0.4364 | 1.54 | 0.1168 | 157760 | 44 |
| 7 | 45461192 | 55025348 | 0.0297 | 0.0312 | 0.5182 | 0.37 | 0.091 | 155663 | 4 |
| 8 | 13442845 | 19170725 | 0.035 | 0.0619 | 0.6053 | 6.08 | 0.2107 | 209024 | 0 |
| 8 | 26904667 | 27684980 | 0.0996 | 0.0168 | 0.6654 | 0.54 | -0.008 | 313103 | 0 |
| 8 | 28217567 | 29212295 | -0.0138 | 0.0222 | 0.7377 | 0.68 | -0.0188 | 194304 | 0 |
| 9 | 194816 | 5233966 | 0.0872 | 0.1489 | 0.571 | 1.31 | 0.1255 | 254345 | 11 |
| 9 | 3624571 | 5069209 | -0.003 | 0.0255 | 0.7702 | 0.46 | 0.2347 | 236930 | 0 |
| 10 | 4970531 | 7454652 | -0.0154 | 0.0181 | 0.5512 | 0.28 | 0.1042 | 157832 | 0 |
| 11 | 42902 | 989171 | -0.0259 | 0.0243 | 0.8601 | 1.6 | 2e-04 | 261982 | 0 |
| 11 | 1717094 | 2887045 | 0.1291 | 0.0295 | 0.7833 | 2.4 | 0.3228 | 232926 | 0 |
| 12 | 3835486 | 14509449 | 0.0352 | 0.0325 | 0.4888 | 1.54 | 0.0831 | 143924 | 0 |
| 12 | 35703715 | 36304361 | -0.0166 | 0.05 | 0.6245 | 0.38 | 0.0546 | 1444439 | 0 |
| 13 | 30006077 | 32432700 | 0.0561 | 0.0774 | 0.6975 | 2.87 | 0.3059 | 219150 | 0 |
| 14 | 13470079 | 14575006 | -0.086 | 0.0208 | 0.761 | 0.71 | 0.1837 | 283673 | 0 |
| 14 | 34884867 | 35895818 | -0.0424 | 0.017 | 0.7657 | 0.3 | 0.0561 | 262086 | 0 |
| 15 | 25408840 | 28073014 | -0.074 | 0.0135 | 0.7139 | 0.72 | 0.1144 | 181994 | 0 |
| 15 | 28507085 | 31697055 | 0.05 | 0.0775 | 0.6882 | 0.14 | 0.0659 | 211765 | 0 |
| 16 | 51475 | 1382176 | 0.0103 | 0.0322 | 0.646 | 0.84 | 0.0247 | 509643 | 0 |
| 16 | 6127996 | 6968254 | -0.0712 | 0.0076 | 0.6481 | 0.26 | 0.0402 | 330029 | 0 |
| 16 | 10128603 | 14367042 | 0.0043 | 0.0207 | 0.4714 | 0.37 | 0.1719 | 195626 | 1 |
| 16 | 34796693 | 35691328 | -0.0892 | 0.0372 | 0.9482 | 0.45 | -0.0203 | 387988 | 0 |
| 17 | 3768696 | 4458439 | 0.1491 | 0.0188 | 0.9046 | 1.69 | 0.6097 | 190618 | 0 |
| 17 | 11894487 | 14516568 | 0.0987 | 0.141 | 0.5793 | 3.72 | 0.0971 | 239485 | 22 |
| 18 | 1680439 | 2232789 | -0.0731 | 0.0104 | 0.8912 | 0.6 | 0.0669 | 674088 | 0 |
| 18 | 32534225 | 34007640 | 0.0477 | 0.0233 | 0.6794 | 0 | 0.0387 | 285507 | 0 |
| 19 | 2765772 | 4034315 | 0.0489 | 0.0918 | 0.7504 | 1.82 | 0.2004 | 189647 | 1 |
| 19 | 24725346 | 25415225 | 0.0812 | 0.1637 | 0.6887 | 3.61 | 0.1061 | 233503 | 7 |
| 20 | 57079 | 5144292 | -0.0058 | 0.036 | 0.5514 | 4.61 | 0.3948 | 238749 | 0 |
| 20 | 14695766 | 18898982 | 0.0438 | 0.0887 | 0.6405 | 2.17 | 0.1082 | 148164 | 2 |
| 20 | 25953482 | 30007504 | 0.0626 | 0.0516 | 0.5095 | 0.14 | 0.1265 | 183742 | 3 |
| 22 | 2331584 | 5114600 | 0.0251 | 0.0253 | 0.4681 | 0.17 | 0.1088 | 374464 | 0 |
| 22 | 30331726 | 32901550 | -3e-04 | 0.0698 | 0.6574 | 0.66 | 0.0585 | 183682 | 3 |
| 23 | 16708893 | 17785992 | 0.0104 | 0.0279 | 0.6202 | 0.51 | 0.0578 | 476754 | 0 |
| 23 | 32884884 | 34749208 | -0.0198 | 0.0512 | 0.5888 | 0.31 | 0.2424 | 189584 | 1 |
| 23 | 43724278 | 44592772 | 0.1167 | 0.0428 | 0.5882 | 0 | 0.0332 | 468786 | 0 |

## Notes

### Competing Interest Statement

The authors have declared no competing interest.

### Summary of Updates

In this version of the manuscript, we expanded the set of parameter used in our simulations, and derived a new measure of recombination landscape evolution: the Population Recombination Divergence Index (PRDI). As a result, we updated Figures 2 and 3, as well as the corresponding text in the Results section. Additionally, we edited the introduction and discussion of our manuscript to more clearly define the scope of the study and strengthen our conclusions.

